# Cytoplasmic circular dsDNA is a key constituent of stress granules

**DOI:** 10.64898/2026.03.12.711345

**Authors:** Natalia A. Demeshkina, Adrian R. Ferré-D’Amaré

**Affiliations:** Laboratory of Nucleic Acids, National Heart, Lung, and Blood Institute, National Institutes of Health, Bethesda, MD, 20892, USA

## Abstract

Stress granules are large cytoplasmic bodies formed in response to environmental insults by eukaryotic cells. Stress granule formation is key for post-stress recovery, and many diseases and infections are characterized by dysregulation of these membraneless organelles. How specific and non-specific macromolecular interactions drive formation of stress granules and other large assemblies is an area of active research. Stress granules are comprised of dense, ∼200 nm cores, and these are known to contain numerous RNAs and proteins. Now, we have discovered that more than half of the nucleic acid content of stress granule cores is circular, double-stranded DNA. We demonstrate cytologically that these extrachromosomal circular DNAs (eccDNAs) colocalize cytoplasmically with canonical stress granule marker proteins, and through CRISPR targeting in yeast, that they are required for stress granule formation upon stress. This discovery thus reveals a key function for eccDNA in the eukaryotic stress response.

## INTRODUCTION

Stress granules are a key cytoprotective response to environmental insults in all eukaryotic cells. Stress granule formation is triggered by various environmental stresses such as chemical exposure, heat shock, oxidative stress and nutrient deprivation^1–6^. Dysfunction of stress granule dynamics is a feature of diverse disease states including cancer^7^ and neurodegeneration^8–11^, and is also a part of the infection mechanism of many viruses^12^. Stress granules arise upon stress-induced reversible stalling of protein synthesis during which most mRNAs are converted from their translationally active state in polysomes into these large cytoplasmic bodies^13–15^. Cytological characterization indicates that stress granules contain dense ∼200 nm cores^16,17^. The biochemical analysis of these cores revealed complex compositions, collectively comprising several thousand candidate proteins and RNAs from diverse cellular pathways. Numerous studies including chemical probing^18–22^ and *in vitro* reconstruction^23–26^ have suggested how some conserved stress granule components interact, and how these interactions may drive stress granule assembly upon stress.

How specific and non-specific macromolecular interactions as well as the physical-chemical properties of biological macromolecules result in the formation of large cytological assemblies such as stress granules and processing bodies in the cytoplasm, and nucleoli, Cajal bodies and speckles in the nucleus is an area of active research. To provide a biochemical counterpart to *in vitro* reconstitution studies, we have previously developed purifications of stress granule cores from *Saccharomyces cerevisiae* and HEK293T cells, and found them to be discrete particles with heterogeneous RNA composition^27^. Now, we have discovered that a large fraction of the nucleic acid content of both yeast and mammalian stress granule cores is DNA, and that this is mostly represented by circular double-stranded molecules. We demonstrate by fluorescence *in situ* hybridization that these circular DNA molecules are genuine components of stress granules in the cytoplasm of human cells. By deploying CRISPR targeting in the cytoplasm, we further show that double-stranded circular DNAs are required for stress granule assembly and that they exist in a chromatin-like state. Our discovery of the functional role of circular DNA in stress granules reveals an unexpected, phylogenetically conserved component of these cytoplasmic membraneless organelles and opens new avenues of research into the fundamental biochemistry of the eukaryotic stress response.

## RESULTS

### Discovery of DNA in stress granule cores

Stress granule cores purified from buddying yeast and mammalian (HEK293T) cells are comprised of complex proteomes^27^. We found that in addition to RNA-binding proteins, the cores are enriched in DNA-binding proteins with functions in chromatin organization and remodeling, such as RFA1 and RAP1 in yeast (Figure 1A), and PARP1 with SMARCC complexes in human (Figure 1B). Other DNA binders previously detected in stress granules^16,27^, including the minichromosome maintenance (MCM) complex and the RUVBL1 and RUVBL2 helicases, are also enriched in our stress granule cores (Figure S1A). Histones had previously been found in stress granule preparations^16,28,29^. Now, our analysis of purified cores shows that their abundance is comparable to those of stress granule marker proteins eIF4A and G3BP1 (Figures 1A and 1B). In the HEK293T cell stress granule cores, the ratio of major histones does not conform to that of canonical nucleosomes, with histones H2B and H4 being most abundant (Figure 1B). In yeast, histone H3 can be epitope-tagged without compromising function^30^. To confirm its unexpected presence in a cytoplasmic membraneless organelle, we subjected stress granule cores purified from a *S. cerevisiae* strain with a chromosome-encoded epitope-tagged H3 to immuno-gold staining and EM (Figure 1C). This revealed robust signal for the histone, consistent with the results of proteomic analyses (Figure 1A).

**Figure 1.**
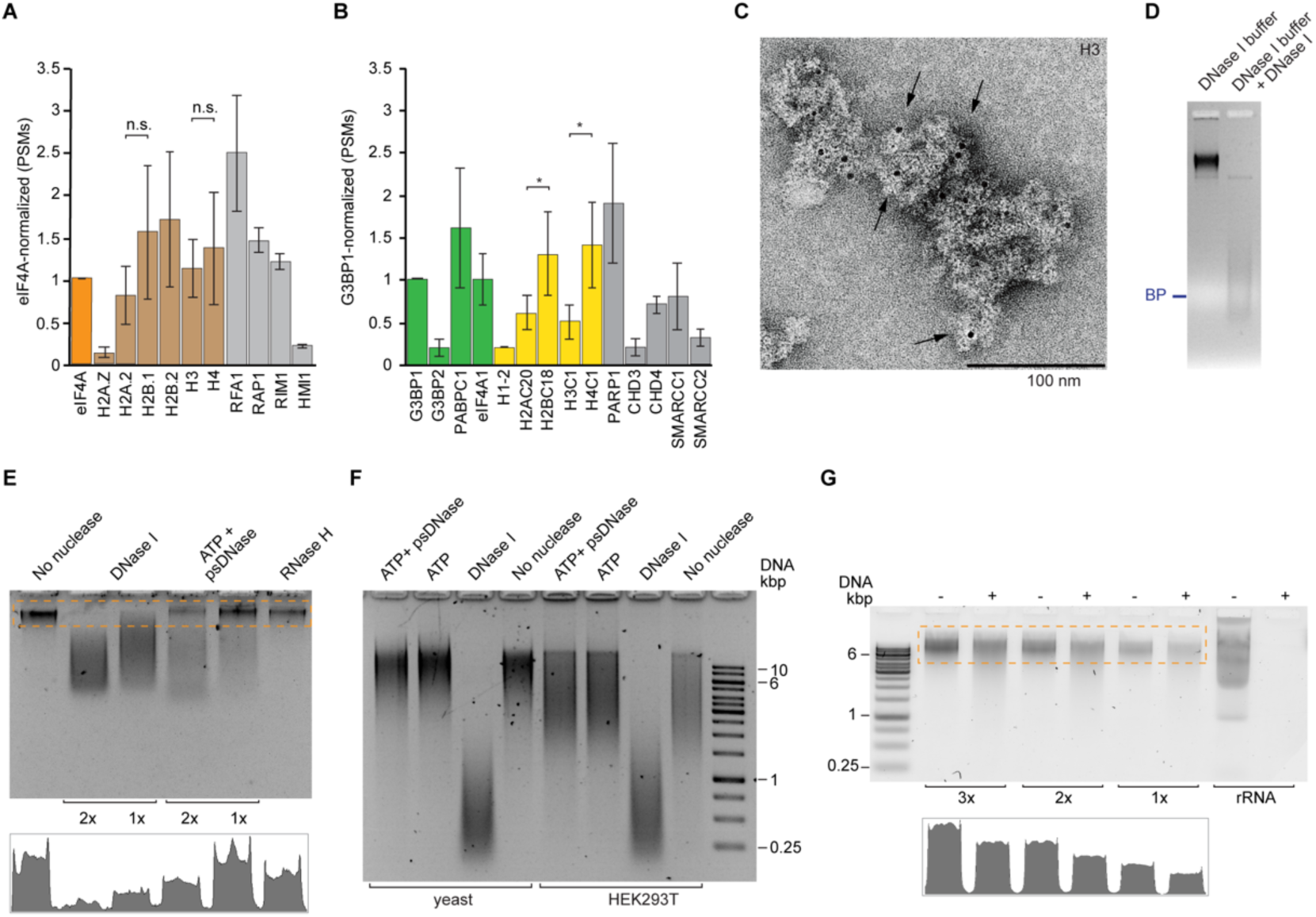
DNA is abundant in yeast and mammalian stress granule cores. (A) Abundance of histones (brown) and other representative DNA-binding proteins (gray) in the proteomes of yeast stress granule cores (strain JD1370 grown in synthetic defined (SD) media), normalized to that of the stress granule marker eIF4A [peptide spectrum matches (PSMs); means from *n* = 2 biological replicates ± s.d.; n.s., no significance; one-tailed Student’s *t*-test]. (B) Abundance of histones and other representative DNA-binding proteins in the proteomes of HEK293T cell stress granule cores, normalized to that of the stress granule marker G3BP1 (PSMs; means from *n* = 3 biological replicates ± s.d.; *, *p* < 0.05; one-tailed Student’s *t*-test. Stress granule markers, histones and non-histone DNA-binding proteins in green, yellow, and gray, respectively. (C) Negative-stain immuno-electron micrograph of stress granule cores from yeast strain YAG1021 carrying a chromosomally encoded FLAG-tagged histone H3. Anti-FLAG antibodies were conjugated to NanoGold beads (arrows; Methods). Magnification (× 23,000). (D) Native agarose gel electrophoresis (SYBR Gold stain) of yeast stress granule cores treated with DNase I. BP, bromophenol blue. (E) DNase I, psDNase and RNase H treatment (the first at two different concentrations) of total nucleic acids extracted from yeast stress granule cores analyzed as in (D). Bottom graph, integrated signal for the dashed box. (F) Comparison of nuclease susceptibility of total nucleic acids extracted from yeast and mammalian stress granule cores, analyzed as in (D). (G) Alkaline hydrolysis of total nucleic acids extracted from yeast stress granule cores (15, 10 and 5 ng) and rRNA (15 ng), analyzed by non-denaturing agarose gel electrophoresis (SYTOX Green stain). Samples marked (+) were incubated with 50 mM KOH for 15 minutes at 95 °C. Bottom graph, integrated signal for the dashed box.

Stress granule cores from yeast and HEK283T cells are heterogeneous particles with median sizes of 135 nm and 220 nm, respectively, that exhibit discrete electrophoretic mobility under non-denaturing conditions^27^. Remarkably, while their electrophoretic behavior is unaffected by treatment with proteinase K or ribonucleases A or T1 under conditions that destabilize other RNPs, such as ribosomal subunits (Figure S1B), treatment with DNase I appears to drastically alter stress granule core stability (Figures 1D and S1C and S1D). To confirm this susceptibility to a deoxyribonuclease, we extracted total nucleic acids from yeast stress granule cores and subjected them to several nucleases (Figure 1E). While DNase I treatment resulted in substantial signal loss, treatment with plasmid-safe DNase (psDNase, an ATPase that does not degrade circular double-stranded DNAs) or RNase H yielded reproducibly modest weakening of the stress granule core nucleic acids band (Figure 1E). The susceptibility to RNase H (an enzyme that targets the RNA strand of DNA:RNA hybrids) suggests that some of the nucleic acids in the particles are heteroduplexes, while the modest signal reduction upon psDNase treatment indicates that a substantial fraction of the DNA is double-stranded and circular. The DNase I susceptibility and resistance to psDNase is phylogenetically conserved, being also a feature of total nucleic acids extracted from HEK293T cell stress granules cores (Figure 1F). These results indicate that circular, double-stranded DNA molecules constitute a major fraction of the DNA content of stress granule cores. To estimate the relative abundance of DNA and RNA in stress granule cores, we subjected total nucleic acids extracted from the yeast particles to harsh alkaline hydrolysis, under conditions where an equivalent amount of rRNA is completely degraded (Figures 1G and S1E). Approximately 50-60% of the stress granule core nucleic acid signal persisted after this treatment. Overall, our analyses revealed that DNA, a large fraction of it circular and double-stranded, is abundant in stress granule cores purified from yeast and mammalian cells.

### Stress granule cores contain eccDNA

Studies over the past sixty years have documented that extrachromosomal circular DNA (eccDNA, originally termed^31,32^ “double minutes”) is widespread in all eukaryotic cells examined, including somatic, germline and cancer cells^33,34^. Advances in DNA sequencing technologies^35,36^ indicate that eccDNAs are polydisperse, ranging from dozens to millions of base-pairs, and that in a given organism, their sequences broadly overlap with the protein coding and non-coding regions of the nuclear genome^37–39^. We adapted the biochemical methodology of Circle-Seq^40^, originally developed for analysis of total eccDNA from yeast cells, and current specialized analytical algorithms (Methods; Figures S2A-S2C). We found that yeast stress granule cores contain abundant DNA circles with characteristics previously identified as hallmarks of eccDNA, namely highly variable size reaching several kilobase-pairs in length (Figures 2A and S2D-S2E), and sequences that map to all sixteen chromosomes (Figure 2B). Characterization of HEK293T cell stress granule core DNA likewise revealed double-stranded circles with a large spread in size and abundance (Figure 2C), and extensive coverage of all autosomes, the X chromosome, as well as the mitochondrial genome (Figures 2D and S2F).

**Figure 2.**
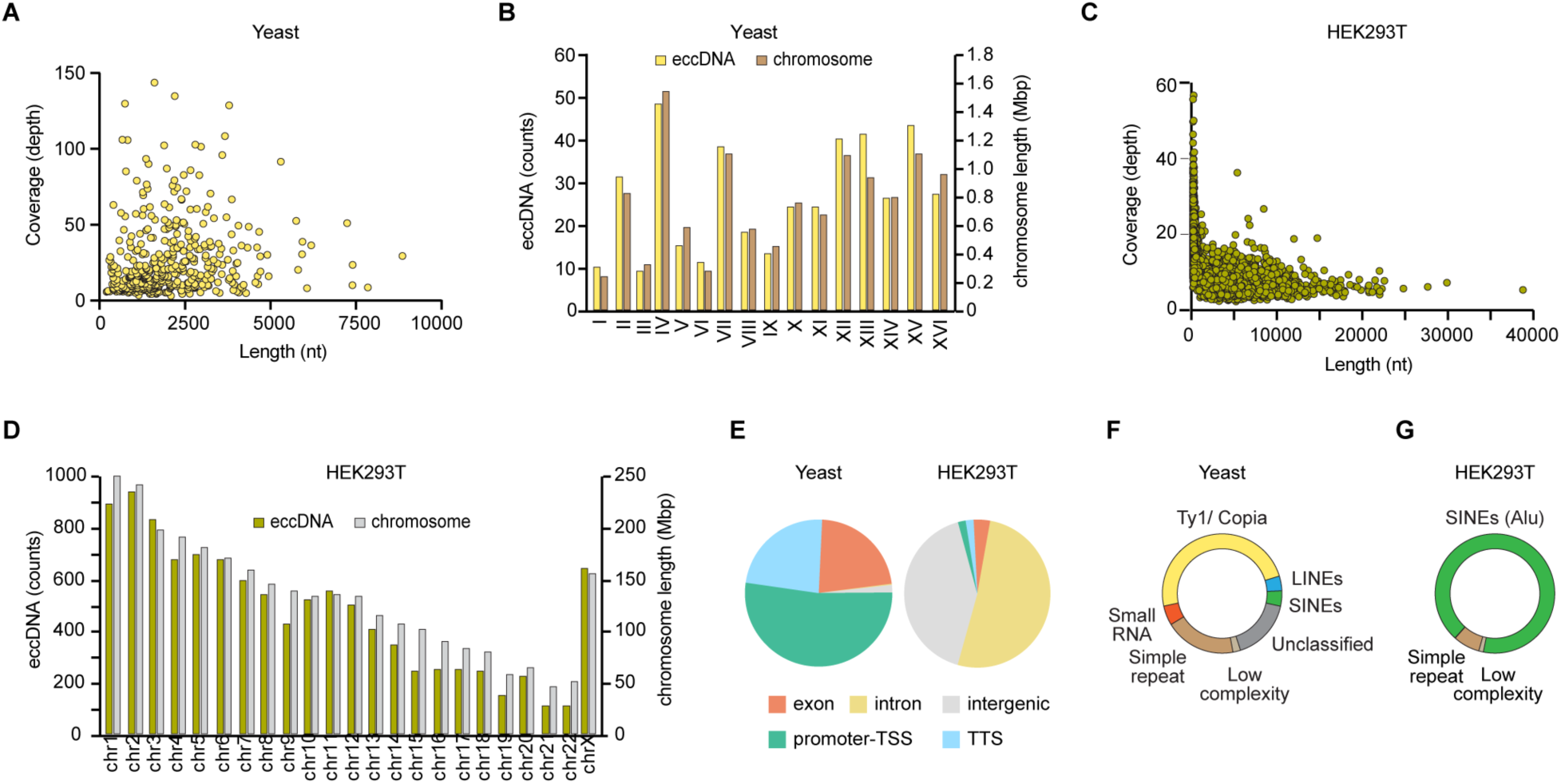
Characteristics of eccDNAs from yeast and mammalian stress granule cores. (A) Abundance and length distribution of high-confidence eccDNAs from yeast stress granule cores (unless noted, the data are from strain JD1370 cultured in SD medium). (B) Genomic coverage, per chromosome, of eccDNAs from yeast stress granule cores. The length of each chromosome is shown for comparison (overall correlation coefficient between relative eccDNA abundance and chromosome length is 0.95). (C) Abundance and length distribution of eccDNA from HEK293T cell stress granule cores. (D) Genomic coverage, per chromosome of eccDNA from HEK293T cell stress granule cores. The length of each chromosome is shown for comparison (overall correlation coefficient between relative eccDNA abundance and chromosome length is 0.95). (E) Relative abundance of gene elements within assemblies of eccDNAs from yeast and human stress granule cores. TSS, transcription start site; TTS, transcription termination site. (F) Repetitive elements within assemblies of eccDNAs of yeast stress granule cores. SINEs and LINEs, short and long interspersed nuclear elements, respectively. (G) Repetitive elements within eccDNAs from human stress granule cores.

The sequences of stress granule core eccDNAs broadly mirror the structure of the nuclear genome. Thus, half of the sequences of yeast stress granule core eccDNAs that map to annotated genes correspond to promoter regions with transcription start sites, and the other half partitions equally between exons and transcription termination sites (Figure 2E). Yeast eccDNA comprise few (2%) intergenic sequences. In contrast, and reflecting the drastically different genomic organization of vertebrates, the eccDNAs from HEK293T cell stress granule cores are comprised mostly of intergenic and intronic sequences, with only 4% exonic regions (Figure 2E). The abundance of repetitive sequences derived from mobile genetic elements has previously been documented in total eccDNAs from a variety of organisms^41–43^. In yeast stress granule core eccDNAs, long terminal repeat (LTR) Ty1-copia retrotransposons are predominant (Figure 2F). These sequences constitute about 2% of the eccDNA assemblies, which is comparable to the coverage^44^ by the LTR elements in the genome of *S. cerevisiae*. For eccDNA from HEK293T cell stress granule cores, the most abundant repetitive sequences are non-LTR retrotransposon SINEs (Figure 2G). These constitute 17% of the eccDNA assemblies, similar to their representation^45^ in the human genome. Overall, our analyses show that the eccDNA complement of yeast and human stress granule cores is broadly similar in structure and sequence to total eccDNAs characterized previously from these sources.

### Stress granules colocalize with eccDNA

Numerous studies have linked eccDNAs to nuclear phenomena, but eccDNA function in the cytoplasm has not been reported. We therefore examined the cellular localization of eccDNAs and DNA-binding proteins present in purified stress granule cores, taking advantage of the suitability of HEK293T cells for light microscopy. Having defined the nuclear boundary by staining against Lamins A and C, we employed hybridization chain reaction fluorescence *in situ* hybridization (HCR-FISH) to detect the coding (*i.e*., sense) DNA strand of the AHNAK gene (Figures S3A-S3B) in cells subjected to oxidative stress (Figure 3A). This clearly demonstrated the presence of AHNAK DNA in both, the perinuclear region and in the cytoplasm, in addition to its expected nuclear localization. The DNA signal for MYC, whose sequence is documented and abundant^46,47^ in mammalian eccDNAs, was likewise present in the cytoplasm and colocalized (Figure 3B) with puncta of the stress granule regulator G3BP and its activator protein Caprin1^23,48^. The DNA signal for AHNAK also colocalized with that for G3BP and Caprin1 (Figure 3C). Both AHNAK and MYC sequences are represented (Figures S3C-S3D) in the stress granule core eccDNAs of arsenite-stressed HEK293T cells, as well as in their transcriptomes^27^. Therefore, we also examined the localization of MYC and AHNAK mRNAs in relation to their cytoplasmic DNA signals (Figures 3D and 3E). After opening DNA strands through heat denaturation (Figures S3B), we found colocalization for a number of RNA foci with the corresponding DNA and G3BP protein signals. However, under these conditions, most G3BP-colocalized DNA and RNA signals were separate from each other. Overall, our observations indicate that stress granule eccDNA can indeed be cytoplasmic, and can colocalize with canonical stress granule marker proteins as well as mRNA.

**Figure 3.**
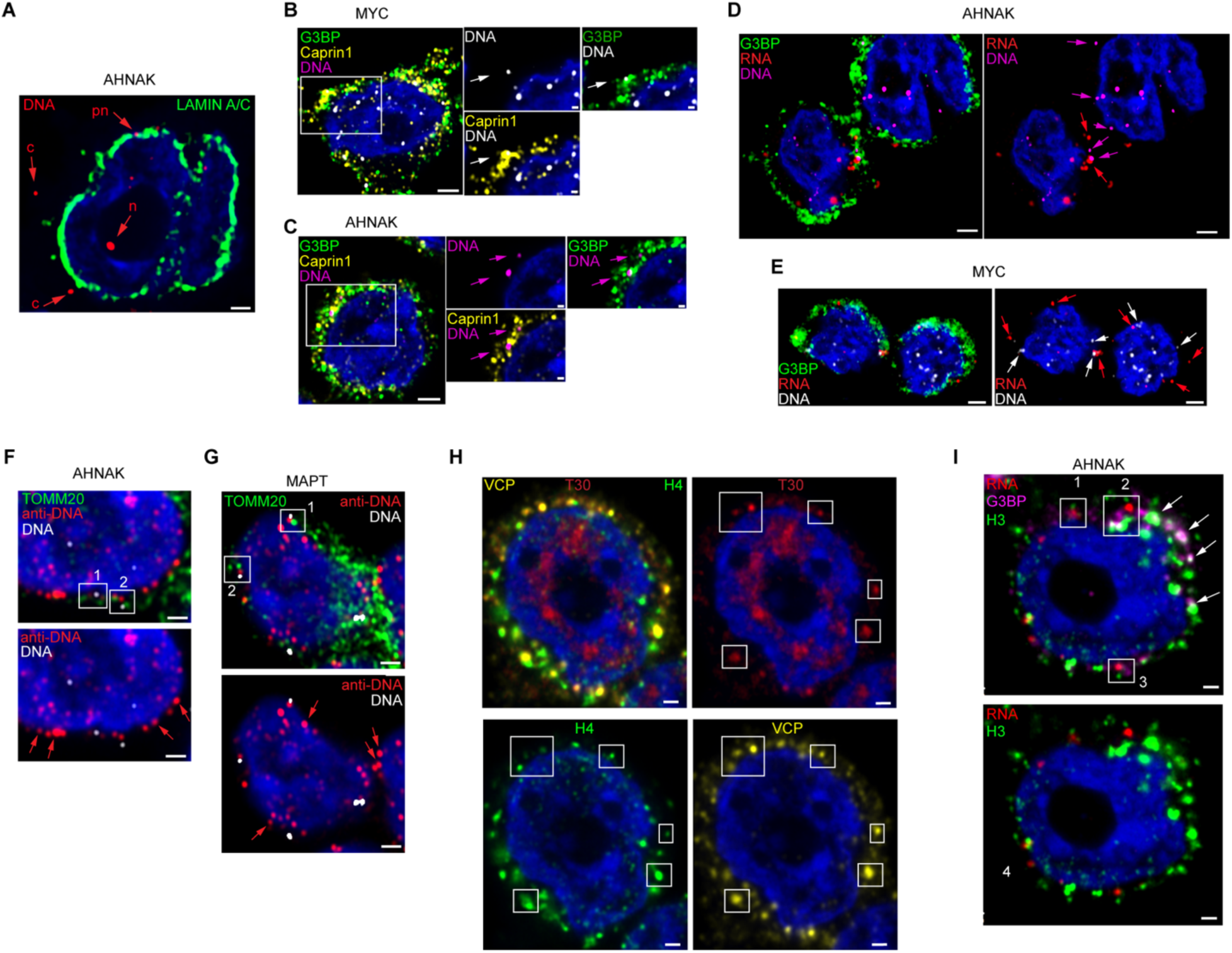
Cytoplasmic eccDNAs colocalize with stress granules in human cells. (A) Partitioning of AHNAK DNA signal between cytoplasm (c), perinuclear (pn) and nuclear (n) volumes of a HEK293T cell. Nuclear envelope is labeled with Lamins A and C. (B-C) Localization of MYC and AHNAK DNA signals in relation to the stress granule marker G3BP and its activator protein Caprin1. (D-E) Concurrent detection of DNA and RNA targets within AHNAK and MYC genes in relation to the stress granule marker G3BP. As in (A-C), denaturing conditions at 75 °C were used to prioritize capturing DNA targets over mRNA targets (red arrows). (F-G) Detection of DNA by conventional anti-DNA IF combined with HCR-FISH against AHNAK and MAPT DNA targets. DNA signals (framed and numbered) proximal to translocase TOMM20 points to relationship with mitochondria. Cytoplasmic DNA without proximal TOMM20 label is marked by red arrows. (H) Colocalization of histone H4 with stress granules turnover protein VCP. Stress granules are demarcated by T30 signal (framed), which represents stress-induced condensation of polyadenylated nucleic acids. (I) Colocalization of histone H3 with the granule marker G3BP and control AHNAK mRNA. Colocalized signals for H3, G3BP, and mRNA are framed and numbered. Arrows point to some H3 and G3BP signals without AHNAK mRNA foci. In all panels, HEK293T cells underwent oxidative stress with 0.5 mM sodium arsenite for 1 hour and 20 minutes at 37 °C, and then were fixed with formaldehyde. Unless otherwise stated, detection of proteins and nucleic acids was carried out by HCR IF and HCR-FISH, respectively. All images represent one confocal Z-stack of 0.2 μm with nuclei stained by Hoechst 33342; proteins and DNA are pseudocolored for clarity. Scale bars 1 μm in (A) and (F-I), 2 μm in (B-E), and 0.5 μm in inserts of (B-C).

To ascertain that the cytoplasmic eccDNA is distinct from the mitochondrial genome, the most studied circular DNA in animal cells, we stained for the translocase TOMM20, widely used to localize mitochondria in cellular imaging studies. We find that a substantial fraction of the cytoplasmic signal for a non-specific anti-DNA antibody does not colocalize with the signal for mitochondria (Figures 3F and 3G). Furthermore, some HCR-FISH signals from probes against the sense strands of AHNAK and MAPT DNA also do not colocalize with mitochondria (Figure 3F and 3G). Thus, the HCR-FISH signal we detect for cytoplasmic eccDNA indicates that these circular molecules are distinct from mitochondrial DNA.

To examine the presence of histones in stress granules, we performed two colocalization experiments with arsenite-stressed HEK293T cells. First, we imaged Valosin-containing protein (VCP), an endoplasmic reticulum AAA+ ATPase that was previously identified as a key player in the dynamics of mammalian stress granules^49,50^. Concurrently, we probed for histone H4 and for polyadenylated RNAs with a T30 probe. This revealed clear cytoplasmic colocalization of punctate VCP signal with those of histone H4 and of T30 (Figures 3H and S3E-S3F). Second, also employing the HCR approach, we imaged histone H3, AHNAK mRNA and G3BP (Figures 3I and S3G-S3H). The AHNAK mRNA has been described^51^ as enriched in stress granules of arsenite-stressed U-2 OS cells. In our imaging experiments, histone H3 colocalized with cytoplasmic G3BP puncta (Figure 3I), and some, but not all such puncta also had clear HCR signal for the AHNAK mRNA. The latter result supports our previous findings^27^ that not every stress granule core contains all mRNAs present in the population of purified particles. Overall, these results are consistent with the abundance of histones in the proteome of stress granule cores purified from HEK293T cells (Figure 1B) and corroborate the presence of these chromatin-component proteins in stress-induced cytoplasmic condensates.

### eccDNA-CRISPR abrogates stress granules

To examine whether the eccDNAs we identified in purified stress granule cores are functional constituents of these membraneless organelles, we repurposed CRISPR technology to target cytoplasmic DNA in *S. cerevisiae* (Figure S4A). For this, we fused *Streptococcus pyogenes* Cas9 to a strong nuclear export signal (NES) and confirmed the cytoplasmic retention of this construct (Cas9_NES_) by imaging its fusion to the fluorescent protein mCherry (Figure S4B). The expression level of Cas9_NES_ was optimized to be moderate (Methods). The CRISPR system was programed with two 20-nucleotide segments from the Ty1 LTR retrotransposon, abundant (Figure 2F) in yeast stress granule eccDNA sequences. The two guide RNAs (gRNAs) were expressed under SNR52 and SUP4 promoters (Figure S4A). The transient expression of the gRNA under SUP4 encoded through a self-cleaving ribozyme-tRNA fusion strategy was verified using a fluorogenic RNA aptamer (see Methods and Figures S4A and S4C). Whereas expression of Cas9 in the nucleus as a nuclear localization signal (NLS) fusion readily triggered formation of stress granules in the absence of any additional stressors (Figure S5A), cytoplasmic expression of the CRISPR components did not (Figures S5B and S5C). This indicates that under these conditions, the programmable, NES-fused endonuclease machinery is not appreciably entering the nucleus.

We then compared the response to oxidative stress of yeast cells expressing an endogenous PAB1-GFP fusion but otherwise wild-type, to that of an isogenic strain expressing the cytoplasmic Cas9_NES_-gRNA holoenzyme. In the absence of stress, diffuse PAB1-GFP signal was present in and indistinguishable between both yeast populations (Figures 4A and 4B). Upon stress, the wild-type yeast cells produced canonical stress granules in the form of large and distinct cytoplasmic puncta (Figure 4A). In dramatic contrast, the isogenic yeast expressing the full CRISPR machinery completely failed to produce stress granules (Figure 4B). We confirmed that CRISPR targeting is operational by isolating stress granule core eccDNA from wild-type yeast, or the corresponding cytoplasmic biochemical fractions from strains expressing either the Cas9_NES_-gRNA holoenzyme, Cas9_NES_ alone, or the holoenzyme assembled with the catalytically dead variant of the endonuclease (dCas9_NES_-gRNA) (Figure S4D). Comparison of the abundance of total eccDNA or of Ty1 DNA sequences specifically show a large reduction only when the active holoenzyme (Cas9_NES_-gRNA) is expressed (Figures 4D and 4E). Thus, cytoplasmic CRISPR targeting of eccDNA suppresses stress granule formation.

**Figure 4.**
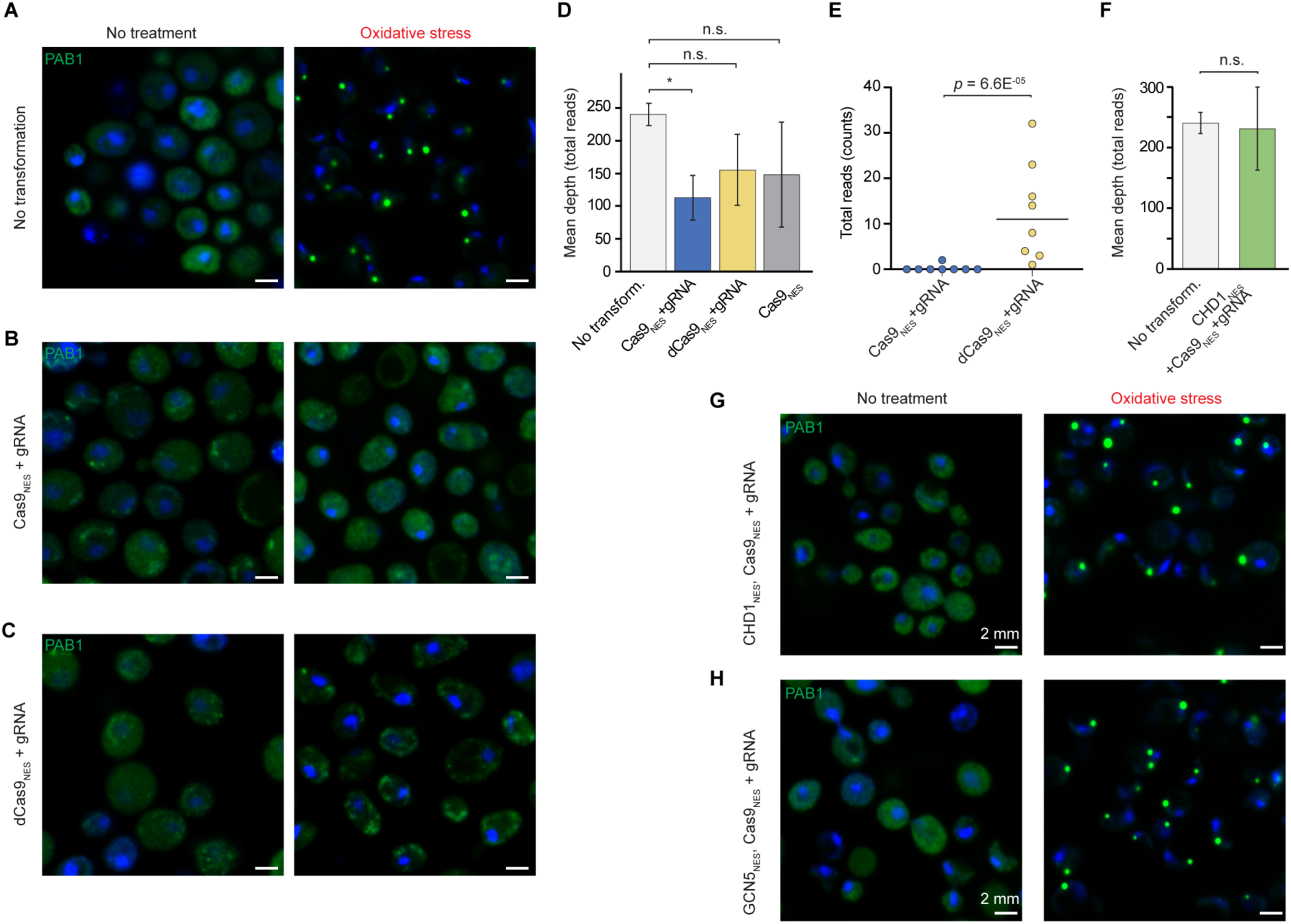
Cytoplasmic eccDNA is required for stress granule formation in yeast. (A-C) Confocal microscopy of *S. cerevisiae* with endogenous PAB1-GFP (green) and transiently expressed (GAL promoter) cytoplasmic CRISPR machinery (Cas9_NES_) with gRNA (Ty1). CRISPR transformant variants are indicated together with treatment conditions. (D) Analysis of total DNA isolated from stress granule cores from wild-type (no transformation) and CRISPR-treated (variants indicated) cells after exposure to oxidative stress with 0.5% w/v sodium azide for 45 minutes (means from *n* = 2 biological replicates ± s.d.). (E) Depletion of 20-nt Ty1 (Ty12^HDV^) targets by enzymatically active cytoplasmic Cas9_NES_ (3rd generation; Methods). Mann-Whitney *U*-test. (F) Effect of cytosolic CHD1_NES_ on abundance of circular double stranded DNA from stress granules isolated from cells co-expressing active CRISPR machinery with gRNA as in (B) (means from *n* = 2 biological replicates ± s.d.). (G-H) Confocal microscopy of untreated and oxidatively stressed *S. cerevisiae* with an endogenous PAB1-GFP fusion (green) and transient (GAL promoter) co-expression of cytoplasmic active CRISPR machinery with cytosolic yeast CHD1_NES_ or GCN5_NES_. In (D) and (F), one-tailed *t*-tests are used; the ordinates are the mean depth of coverage over sixteen chromosomes (mitochondrial genome is omitted); n.s., no significance; *, *p* < 0.05. All images represent one middle confocal Z-stack of 0.14 μm with nuclei stained by DAPI (blue). Scale bar 2 mm.

### Chromatin factors suppress CRISPR effect

Unexpectedly, cytoplasmic expression of the catalytically inactive dCas9_NES_-gRNA holoenzyme targeting Ty1 resulted in abrogation of stress granule formation that was comparable to the effect of cytoplasmic expression of the catalytically active Cas9_NES_-gRNA counterpart (Figures 4B and 4C and S4A). Indeed, expression of Cas9_NES_ together with a gRNA targeting it to a sequence that is absent in the yeast genome was sufficient to abolish stress granule formation (data not shown). These results suggest that a biochemical activity of the CRISPR system distinct from site-specific, gRNA-targeted endonucleolytic DNA cleavage is responsible for the observed abrogation of stress granule formation. Previous biophysical characterization of Cas9-gRNA shows that the holoenzyme (independent of its ability to cleave DNA) associates strongly with DNA, and efficiently performs a one-dimensional diffusive target search^52–54^, which contributes to its transcriptional repression properties^55^. We therefore hypothesized that non-specific cytoplasmic association of the CRISPR machinery with DNA is disrupting molecular interactions of eccDNA that are required for stress granule formation.

The abundant histones in purified stress granule cores (Figures 1A and 1B) and their colocalization with several canonical stress granule markers (Figure 3) suggest that the eccDNAs that form part of these cytoplasmic bodies may be organized in a manner that resembles nuclear chromatin. We reasoned that cytoplasmic expression of proteins that associate with chromatin may be sufficient to compete with the binding, or block the one-dimensional diffusion of the CRISPR machinery on cytoplasmic eccDNAs, thereby restoring the ability of cells to form stress granules. We tested this by expressing, separately, two structurally and biochemically unrelated chromatin interacting enzymes, the chromatin remodeler CHD1, and the histone acetyltransferase GCN5, both as NES fusions (Figure S6A). We first determined that expression of the CHD1_NES_ fusion fully overcomes the reduction in cytoplasmic eccDNA (Figure 4D) that results from cytoplasmic expression of the CRISPR machinery (Figure 4F), suggesting that the chromatin remodeler effectively competes with (catalytically active) Cas9_NES_-gRNA for access to the cytoplasmic eccDNA. Examination of the response to oxidative stress of yeast cells expressing either CHD1_NES_ or GCN5_NES_ in addition to catalytically active Cas9_NES_-gRNA revealed that either chromatin-interacting protein was sufficient to restore stress granule formation to an extent indistinguishable from wild-type cells (Figures 4G and 4H). This pronounced effect was not elicited by the expression vectors alone (Figure S6). An identical response was obtained when the two chromatin-targeted enzymes were expressed in cells expressing the catalytically inactive CRISPR machinery (dCas9_NES_-gRNA, not shown). These results strongly suggest that cytoplasmic eccDNA is packaged in a chromatin-like state, and that interactions that it needs to make with other cellular components in order to drive stress granule formation are disrupted by the tight DNA binding or one-dimensional diffusion of the CRISPR holoenzyme.

### eccDNA-CRISPR impairs hypoxia recovery

In a variety of contexts, it has been observed^21,56^ that the ability of cells to make stress granules correlates with their successful recovery upon cessation of the noxious conditions. To determine if the ability of cytoplasmic Cas9_NES_-gRNA machinery to suppress stress granules is functionally suppressing the stress-protective biological role of these condensates (and not just their cytological appearance), we assayed yeast expressing various components of the CRISPR system for their ability to recover from sodium azide-induced stress. When cells stressed in solution were spotted on a recovery (non-stress) solid medium (Figure 5A), their growth largely correlated with their ability to make stress granules (Figures 4 and S4). Analysis of colony numbers (Figure 5B) demonstrates that the most efficient recovery occurred in yeast expressing just Cas9_NES_ or gRNAs, neither of which appreciably suppress stress granule formation. Under our experimental conditions, co-expression of CHD1_NES_ with enzymatically active or dead Cas9_NES_ CRISPR holoenzymes had a strong protective effect (Figures 5A and 5B). We further assayed recovery in liquid cultures, finding a corresponding pattern of delayed growth for the strains expressing CRISPR machinery that suppresses stress granule formation (Figure 5C). Thus, differential recovery from hypoxia provides evidence of the requirement for eccDNA accessibility for cellular protection from stress that correlates precisely with the cytologically determined ability to form stress granules.

**Figure 5.**
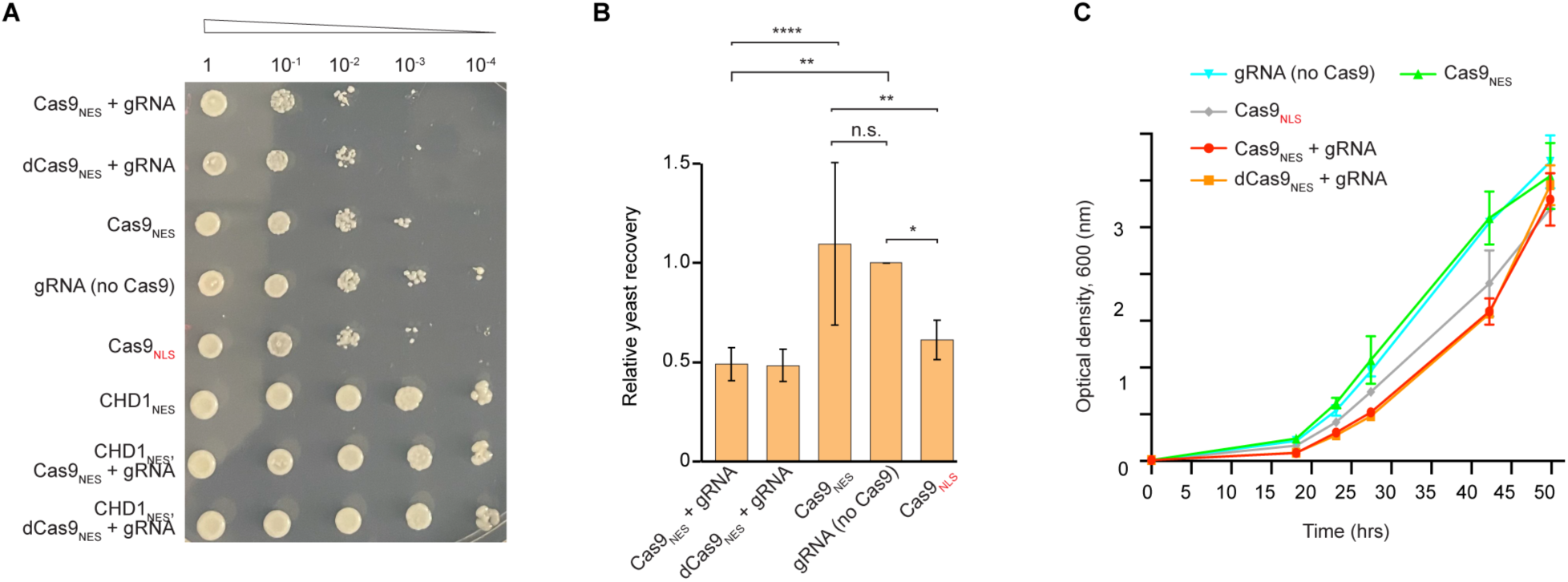
CRISPR-mediated suppression of stress granules compromises recovery from hypoxic stress. (A) Recovery of CRISPR transformants after exposure to oxidative stress (45 minutes at 30 °C) applied in the early log phase (Methods). Spotting assay was carried out under moderate Cas9_NES_ induction on SD(-URA) solid medium with D(+)-glucose (1.5% w/v) and D(+)-galactose (0.75% w/v) using indicated ten-fold serial dilutions. Stress-induced phenotypes of the CRISPR variants with controls are presented in Figures 4 and S5. (B) Quantitation of post-stress recovery efficiency as in (A) for indicated CRISPR transformants. Spotting assays were carried out on solid medium at 30 °C for 2.5 days as in (A) and quantified using the second (10^-2^) dilution. Data are mean for *n* = 3 ± s.d. and 44 technical replicates. Data were normalized to values of gRNAs alone; n.s., no significance; *, *p* < 0.05; **, *p* < 0.01; ****, *p* < 0.0001 (one-way ANOVA followed by Tukey’s test with a 95% confidence interval). (C) Representative time-course for post-stress recovery of CRISPR transformants in the SD(-URA) liquid medium as in (A) (*n* = 2 ± s.d.; technical replicates). The growth was initiated with the second (10^-2^) dilution.

## DISCUSSION

By analyzing purified stress granule cores^27^ from yeast and mammalian cells, we have discovered that circular, double-stranded DNA is a major component of these cytoplasmic bodies. Hitherto, it has been generally thought that stress granules are comprised exclusively of proteins and RNA, but some scattered observations have hinted at the presence of DNA. Thus, an imaging study of human osteosarcoma (U-2 OS) cells reported that treatment with a DNase resulted in substantial loss of the stress granule signal^23^. A pioneering biochemical analysis noted that stress granule fractions enriched from yeast and U-2 OS cells were resistant to ribonucleases^16^, observations that we have now extended to purified stress granule cores (Figure S1B). A medicinal chemistry study that investigated the binding of G-quadruplex-specific ligands to stress granules^57^ noted the persistence of binding of these compounds to the granules even after RNase treatment, and proposed that those stress granules contain DNA. Our biochemical analysis indicates that more than half of the nucleic acid content of purified yeast and mammalian stress granule cores is DNA, and that most of the DNA is double-stranded and circular (Figures 1E and 1F). Circular DNA molecules would not have been detected in stress granules by conventional sequencing approaches, which rely on defined elements at the ends of linear molecules, or lost by shearing steps prior to template preparation.

Since their discovery sixty years ago^31,32^, eccDNAs have been implicated in diverse biological phenomena in eukaryotes. In particular, it has been shown that large extrachromosomal DNA circles that encode oncogenes and regulatory elements are abundant in various malignant tumors^34,58,59^, and may be not just a mechanism for amplification for oncogenes, but could also play a role in immunomodulation and inflammation^60^. A cytoplasmic function for eccDNAs has hitherto not been described. In vertebrates, the cyclic GMP–AMP synthase (cGAS)–stimulator of interferon genes (STING) pathway recognizes damaged cytoplasmic self-DNA and foreign (*e.g*., viral) double-stranded DNA, and induces the innate immune response^61^. This pathway can be also experimentally triggered by the transfection of protein-free plasmids. Chromatinization of nuclear DNA precludes activation of cGAS^62^. Our observations that cytoplasmic expression of the CRISPR machinery suppresses the ability of cells to form stress granules, and that this suppression can be relieved by cytoplasmic expression of chromatin-interacting proteins strongly suggest that the eccDNA component of stress granule cores is packaged in a manner that resembles nuclear chromatin. This is consistent with abundance of histones and other DNA-associated proteins in the proteomes of stress granules^16,27–29^. How eccDNAs are chromatinized and exported from the nucleus is an important future area of investigation.

Our determination of the cytoplasmic colocalization of eccDNAs with stress granules (Figure 3), and of the suppression of stress granule formation *in vivo* by non-endonucleolytic activities of the Cas9-gRNA holoenzyme (Figure 4) suggest that the circular DNA molecules may function in nucleating stress granule assembly. In this regard, our observation that the RNA component of stress granule cores is susceptible to RNase H (Figure 1E) suggests that DNA-RNA heteroduplexes are present in stress granules. Because eccDNAs broadly cover the nuclear genome (Figure 2), in principle, any element of the transcriptome will be sequence-complementary to some eccDNA. Several recent studies suggest that stress granules nucleate from “seeds” that exist under stress-free conditions, and that these form the basis for the assembly of the mesoscopic membraneless organelles that become apparent under stress^18,29,63–65^. It is possible that eccDNAs participate in formation of such seeds. Investigating the abundance, composition and interactome of cytoplasmic eccDNA in the absence of stress will, therefore, be of great interest. Recent developments suggest that long non-coding RNA (lncRNA) molecules play key nuclear roles modulating chromatin structure and participating, by binding to DNA, in the activity of transcriptional enhancers and their coalescence into condensates^66,67^. As lncRNAs are abundant in human stress granule cores, they may play analogous roles in the cytoplasm by associating with eccDNA. Collectively, our work reveals an unexpected linkage between the nuclear genome and the eukaryotic stress response, and suggests that DNA-RNA interactions may play fundamental biological roles in the cytoplasmic compartment.

## ACKNOWLEDGEMENTS

We are grateful to Y. He from the fermentation facility (NHLBI) for HEK293T cell cultures; to X. Wu of the Light Microscopy Core (NHLBI) for confocal microscopy support; the staff of the Electron Microscopy Core (NHLBI) for technical assistance; and T. Dever (National Institute of Child Health and Human Development), H. Shroff (Janelia Research Campus), T. Tsukiyama (Fred Hutchinson Cancer Research Center), and C. Jones and all other members of the Ferré-D’Amaré laboratory for discussions. We acknowledge A. Cait, H. Hassan, and E. Scott of Bridge Informatics (Cambridge, MA) for computational analyses and assembly of circular DNA. The contributions of the NIH authors were made as part of their official duties as NIH federal employees, are in compliance with agency policy requirements, and are considered Works of the United States Government. However, the findings and conclusions presented in this paper are those of the authors and do not necessarily reflect the views of the NIH or the U.S. Department of Health and Human Services.

## AUTHOR CONTRIBUTIONS

Conceptualization, N.A.D. and A.R.F.; methodology, N.A.D.; investigation and formal analysis, N.A.D. and A.R.F; discussion and writing, N.A.D. and A.R.F.

## METHODS

### Materials

The yeast strain with a chromosomally encoded PAB1-GFP fusion was from the Yeast GFP Clone Collection (Thermo Fisher Scientific). Strains JD1370 (*MATalpha trp1 ura3 leu2 [L-A] [Mi] PEP4::HIS3 NUC1::LEU2*), and YAG1021 (*MATalpha W303 hht2::FLAG-HHT2-URA3*), were kindly provided by Professors J. Dinman (University of Maryland, College Park, MD, USA) and A. Gunjan (Florida State University, Tallahassee, FL, USA), respectively. All plasmids employed for the CRISPR experiments were generated based on the pML104 backbone (Addgene #67638). The pBY011 plasmids encoding *S. cerevisiae* CHD1 and GCN5 genes were purchased from the DNASU (clone IDs: ScCD00095307 and ScCD00011312, respectively). Several alterations of pBY011 and pML104 were performed by GenScript.

### Stress granule cores

Purification of stress granule cores was performed as described^27^. For yeast, cells were grown in SD medium with uracil (20 mg/l) (wild-type) or without uracil (CRISPR transformants). For each CRISPR SDGC-SEC-purified stress granule cores, fermentation was 2-3 l.

### Electron microscopy

For localizing FLAG-tagged histone H3, purified yeast stress granule cores (strain YAG1021) were first embedded on glow-discharged grids (50 s at 21 °C) (Electron Microscopy Sciences, FCF300-CU Formvar/Carbon Mesh, Copper). The grids were treated with 0.1% CWFS gelatin (Aurion), monoclonal Anti-FLAG M2 antibodies (Sigma-Aldrich, Cat# F1804) and secondary antibodies (1.3 nm Nanogold-IgG Goat anti-mouse IgG) (Nanoprobes, Cat# 2001) as described^27^.

The grids were washed seven times with 1X TBS for 1 min. The grids were then gold-enhanced according to the vendor’s protocol (Nanoprobes, GoldEnhance EM plus) for 15-20 s, washed eleven times for 1 min in distilled water and stained with 1% (w/v) uranyl formate (Electron Microscopy Sciences, Cat# 22450) for 30 s; the staining was repeated once more. Micrographs were acquired on FEI Tecnai T12 with a focus on the electron dense gold particles.

### Treatment of stress granule cores with proteinase K and nucleases

Yeast stress granule cores (∼ 0.01 A_260_ units) or 60S subunits (0.07 A_260_ units) were incubated with 30 μg of proteinase K (Invitrogen, Cat# 25530-049), RNase A/T1 mix (1 unit of RNase A and 40 units of RNase T1) (Invitrogen, Cat# AM2286) or a mixture of the three enzymes for 1 day at 37 °C in the final volume of 25 μl. Before digestion, the reaction mixtures were supplemented with reaction buffer M [for proteinase K: 30 mM Hepes-KOH pH 7.5 (at 20 °C), 5 mM Mg(OAc)_2_, 125 mM KCl, 0.25 mM EDTA-KOH, 8.5% (w/v) D-Mannitol ] or buffer ‘A/T1’ [for A/T1 or A/T1+K: 10 mM Tris-HCl 7.5 (at 20 °C), 300 mM NaCl, 5 mM EDTA-KOH]. In parallel two controls of the cores and 60S were incubated in buffer M on ice or at 37 °C. The reaction mixtures (7 μl) were complemented with the loading buffer (3 μl of 1X TAE , 0.1% (w/v) BP, 30% glycerol (v/v), 34% (w/v) sucrose) and assayed by agarose gel electrophoresis [1% (w/v), 1X TAE]. The gel was stained by ethidium bromide in 1X TAE. Separately, the same amount of yeast stress granule cores or 60S and corresponding controls were incubated with or without TURBO DNase (0.07 units/ml) (Invitrogen, Cat# AM2238) in the enzyme reaction buffer (1X) for 1 h at 37 °C in 25 μl. The assay was as above.

### Extraction of total nucleic acids from stress granule cores

Stress granule cores (0.2-0.5 A_260_ units) were incubated with 400 μg of proteinase K for 16 h at 37 °C. Total nucleic acids were extracted using phenol:chloroform and phase lock tubes (Quantabio, 5PRIME, Heavy, Cat# 2302830) followed by MicroSpin S-400 HR column elution (Cytiva, Cat# 27514001). The extraction was repeated once. Total extracted nucleic acids were precipitated in 70% (v/v) ethanol and 300 mM sodium acetate (pH 5.2) in the presence of GlycoBlue carrier (Invitrogen, Cat# AM9516) at -20 °C, overnight. The samples were centrifuged at 12,000 x *g* at +4 °C for 20 min and the pellets were washed with 70% (v/v) ethanol, recentrifuged, washed again, air-dried at room temperature for 20-25 min and dissolved in 10 mM Tris-HCl, pH 8.0.

### Treatment of total nucleic acids with nucleases

Total nucleic acids from yeast stress granule cores (125 ng per 100 μl) from yeast (strain JD1370 grown in SD; Figure 1E) were incubated with 0.001 or 0.003 units of TURBO DNase, 0.2 or 0.4 units of psDNase (Biosearch Technologies, Cat# E3101K) with 1 mM ATP or with 9 units of RNase H (New England Biolabs, Cat# M0297). Each reaction mixture was supplemented with the corresponding enzyme reaction buffer (1X) and incubated for 40 min at 37 °C. The reactions (10-15 μl) were mixed with the loading buffer (see above) and assayed by agarose gel electrophoresis [1% (w/v), 1X TAE]. The electrophoretic run was stopped when the BP blue dye migrated 1 cm from the loading wells; the gel was stained with SYBR Gold (Invitrogen, Cat# S11494). Intensities of major bands (Figure 1E) were analyzed using Fiji (ImageJ) (version 2.9.0/1.54p). Alternatively, total nucleic acids from stress granule cores (375 ng per 100 μl) from yeast (strain JD1370 grown in SD; Figure 1F) and HEK293T cells (early log phase; Figure 1F) were treated with 0.002 units of TURBO DNase, 1 mM ATP or 0.075 units of psDNase with 1 mM ATP in the corresponding enzyme reaction buffers (1X) for 30 min at 37 °C. The reaction mixtures were assayed as above with BP blue dye at 3.3 cm from the loading wells. Staining was with SYBR Gold.

### Alkaline hydrolysis of total nucleic acids

Alkaline hydrolysis of total nucleic acids (1.2 μg) extracted from yeast stress granule cores was performed by heating samples in 9.5 μl of 53 mM KOH for 15 min at 95 °C using a C1000 Thermal Cycler (BioRad). Next, 8 μl of the samples were diluted with 160 μl of ultra-pure water to bring KOH to 2.5 mM. The samples were further diluted with 2.5 mM HCl and 2.5 mM KOH to adjust pH to 8.0, mixed with the loading buffer and assayed by agarose gel electrophoresis as above. Various quantities of the same treated sample (15, 10 and 5 ng in Figures 1G and S1E) were analyzed in parallel with the matching quantities of the control total nucleic acids, which were not treated with KOH and were diluted as above with ice-cold ultra-pure water. Total ribosomal RNA (rRNA; 15 ng) extracted as above from pellets of the sucrose density gradient centrifugation step^27^ applied to HEK293T cells was used as a positive control. The agarose gel was stained by SYTOX Green (Invitrogen, Cat# S7020) in 1X TAE for 15 min at 21 °C and imaged using Typhoon scanner (Amersham, software 2.0.0.6) and Cy2 channel. The intensities of major bands were analyzed using Fiji (ImageJ) (version 2.9.0/1.54p). Alternatively, the same reaction mixtures were analyzed with additional control of plasmid DNA (15 ng; Figure S1E) and the gel was stained by SYBR Gold to confirm the results on relative abundance of DNA and RNA in the cores.

### Enrichment and amplification of double-stranded circular DNA

Nucleic acids enriched in DNA were extracted from yeast stress granule cores (0.3-0.5 A_260_ units; strain JD1370 grown in SD medium) using the TRIzol reagent (Invitrogen, Cat# 10296010) by recovering material from the interphase and further purifying it using DNA Clean & Concentrator-25 (DCC-25) kit (Zymo Research, Cat# D4034). The samples were further treated with RNase A/T1 mix (see above) for 1 h at 37 °C and purified again using DCC-25 minicolumns. For each sample, 1.5 μg of nucleic acids enriched in DNA were digested with 60 units of psDNase (Biosearch Technologies, Cat# E3101K) and 1 mM ATP in psDNase reaction buffer at 37 °C. The digestion was carried out for additional 4 days with the fresh enzyme and ATP supplemented every 24 h (in total 60 units of psDNase were added per 1.5 μg). To eliminate remaining RNA, 5 μl of psDNase-treated sample were incubated with 5 units of RNase H (New England Biolabs, Cat# M0297) in total volume of 50 μl for 40 min at 37 °C in RNase H reaction buffer. Further, 5 μl of each RNase H reaction mixture was taken for DNA amplification using phi29 and random primers (Qiagen REPLI-g kit, Cat# 150043) at 16 °C overnight. The phi29 enzyme was inactivated at 65 °C for 10 min and the DNA was purified using DCC-25 minicolumns as above. Samples were sequenced by SeqCenter through a hybrid sequencing approach using Illumina (paired short reads) and Oxford Nanopore technology (long reads; ONT) platforms (Medium Combo package; accessions: SRR35009460, SRR35009461, SRR35009462, SRR35009463). Illumina reads provided high base accuracy, while ONT reads allowed for resolution of larger, repetitive, or complex structures. The sequencing analysis including assemblies into candidate DNA circles was performed by Bridge Informatics.

DNA from yeast modified by CRISPR machinery was extracted similarly to the method described above. Firstly, the samples (0.3-0.5 A_260_ units) were digested with RNase A/T1 mix (Invitrogen, Cat# AM2286) for 1 h at 37°C, followed by digestion with proteinase K (Invitrogen, Cat# 25530-049) for 1 h at 37°C at 450 rpm with addition of SDS to 1% (w/v). DNA was further extracted using TRIzol and cleaned up using DCC-25 as above. The samples were further incubated with exonuclease VIII (20-30 units per 1 μg of DNA) (New England Biolabs, Cat# M0545) up to 15 h at 37 °C. Exonuclease T (20 units per μg of DNA) (New England Biolabs, Cat# M0265) was then added, and samples were further incubated for 16-17 h at 25 °C. Reactions were quenched by heating at 70 °C for 30 min, and DNA was cleaned using DCC-25. Next, DNA was treated with psDNase in the presence of 1 mM ATP and 1 mM DTT as above using ten-fold excess of the enzyme over recommended enzyme quantity during every new daily addition over 5 days. Finally, 5 μl of each reaction mixture was used for phi29 amplification as above; the DNA was purified using DCC-25 minicolumns. The initial analysis was performed by Genewiz (Azenta Life Sciences) using the Illumina MiSeq 2 × 250 bp paired-end sequencing platform. After quality control, reads were trimmed, and low-quality bases were removed (Trimmomatic 0.36). Reads were mapped to the S288C reference genome using the BWA package (0.7.12). Mapping to multiple regions of the genome was allowed to collect information about repetitive regions. The final analysis of chromosomal (cumulative: I-XVI) coverage was carried out with subtraction of regions for URA3 (chr V: 116091-117045), the GAL1 promoter (chr II: 278565 – 279010), rDNA (chr XII: 451426-468907) and RPL4B (chr IV: 471853 – 472941) (Samtools (v.1.17), Bedtools (v.2.30.0); properly paired reads only). Analysis of mean depth of coverage (cumulative: I-XVI) was performed with subtraction of regions for URA3, GAL1, rDNA as above and CHD1 (chr V: 505392 - 509798) (all reads considered).

Extraction of DNA from stress granule cores purified from HEK293T cells stressed at early and late log phases was performed as follows. Firstly, HEK293T stress granule cores (0.2-0.3 A_260_ units) were incubated with 400 μg of proteinase K (Invitrogen, Cat# 25530-049) for 16 h at 37 °C. Then nucleic acids were purified by 2x phenol:chloroform using phase lock tubes (Quantabio, 5PRIME, Heavy, Cat#2302830) and MicroSpin S-400 HR columns (Cytiva, Cat. No. 27514001). Nucleic acids were precipitated with 3 volumes of ethanol in the presence of 300 mM Na-acetate (pH 5.2) and GlycoBlue carrier (Invitrogen, Cat# AM9516). Total nucleic acids (1.5 μg) were then subjected to digestion with psDNase (3 units) in the presence of 1 mM ATP in the psDNase reaction buffer for 1 day at 37 °C. The reaction mixtures were supplemented with fresh psDNase (1.5 unit), 1 mM ATP and 1 mM DTT and incubated for additional 2 days at 37 °C. Each reaction mixture (27 μl) was further diluted 1.5 times to adjust Tris-HCl (pH 8.0) and MgCl_2_ to 25 mM and 10 mM, respectively. RNase A (1.2 μg) (Thermo Scientific, Cat # EN0531) was then added and digestion was carried out for 1 h at 37 °C. Next, 5 units of thermostable RNase H (New England Biolabs, Cat# M0523) was added and the reaction was incubated for 30 min at 86 °C in the RNase H reaction buffer. Prior to the amplification step, samples were incubated with LiCl added to 200 mM for 10 min at 37 °C. The amplification with phi29 was performed as above using 5 μl of the LiCl-treated samples. Amplified DNA was then debranched with endonuclease T7 (New England Biolabs, Cat# M0302) for 1 h at 37 °C. The T7 reaction mixtures were cleaned using 1x AMPure XP magnetic beads (Beckman Coulter, Cat# A63880). Alternatively, DNA was amplified from total nucleic acids as above without treatment with RNase A and LiCl, and the debranching step was also omitted (accessions: SRR35178673, SRR35178674, SRR35178675, SRR35178676, SRR35178677, SRR35178678). The samples were sequenced by SeqCenter using Illumina and ONT platforms (Large Combo). Assemblies into candidate DNA circles were performed by Bridge Informatics.

### Assembly of DNA reads into candidate circles

#### Read processing and quality control

Illumina paired-end reads were processed in the Circle-Map pipeline^68^ starting with bwa mem (v0.7.17) alignments to either the GRCh38 reference genome or *S. cerevisiae* R64 with default-q trimming, followed by coordinate (samtools sort) and query-name sorting, header inspection (samtools view -H), and header correction (samtools reheader). Reads were then deduplicated with fastq-uniq from BBMap (v38.96) after adapter and low-quality base removal using fastp (v0.23.4, default adapter detection, Q20 cutoff, min length 30 bp). Long reads obtained via ONT were evaluated with the CReSIL^69^ NanoPlot module (–maxlength 40000–plots dot–legacy hex) and mapped to reference genomes using minimap2 (v2.24) with ONT-specific parameters-ax map-ont --secondary=no --split-prefix chimeric_splits -z 400,50 -p 0.2 --max-chain-skip 25 -t 16. The ONT reads were additionally assessed for ghost or shadow sequences to minimize spurious eccDNA assemblies. All retained reads were converted to FASTA or multi-FASTA format for downstream analysis.

#### High-confidence eccDNA assembly

High-confidence eccDNA assemblies were generated by integrating Illumina and ONT datasets. For Illumina data, Circle-Map (v1.1.4) was used in two stages: ReadExtractor to isolate circular read candidates from re-headed BAMs, followed by Realign against the reference genome using both coordinate-sorted and query-name-sorted BAMs. For yeast (accessions: SRR35009460, SRR35009461, SRR35009462, SRR35009463) and some human (accessions: SRR35178673, SRR35178674, SRR35178675, SRR35178676, SRR35178677, SRR35178678) datasets sequenced via the hybrid approach assemblies of high-confidence eccDNAs were defined using *overlap* (Figure S2C), in which Circle-Map and CReSIL.bed outputs were intersected with bedtools intersect -wa -wb to retain only coordinates detected by both platforms. For the second part of human data (accessions: SRR34981492, SRR34981493, SRR34981494, SRR34981495, SRR34981496, SRR34981497, SRR35328250, SRR35328251, SRR35328252, SRR35328253, SRR35328254, SRR35328255), due to low Illumina coverage and the predominance of ONT calls, a *merge* approach was used (Figure S2C). In this case, Circle-Map and CReSIL.bed outputs were combined with bedtools merge, retaining the union of intervals and prioritizing Circle-Map scores when both were present.

#### Length distribution and coverage analysis

Length distribution of eccDNA assemblies was quantified by binning assembled sequences into defined size categories. Coverage profiles were generated from sorted BAM files using bedtools genomecov and converted to bigWig using UCSC bedGraphToBigWig (v4) with chromosome sizes derived from samtools faidx. For yeast high-confidence assemblies, coverage was calculated relative to the *S. cerevisiae* R64 genome; for human merged assemblies, coverage was calculated across the full GRCh38 genome assembly.

### Genome annotation of coding and non-coding elements for yeast

All high-confidence (overlap) eccDNA coordinates were annotated against *S. cerevisiae* R64 reference genome using HOMER (v4.11) (annotatePeaks.pl). Each eccDNA was assigned a feature category based on its overlap with gene models. Coding regions included full or partial overlap with exons and protein-coding genes. Non-coding regions encompassed promoters, untranslated regions (UTRs), introns, and intergenic regions. The representation of coding versus non-coding eccDNAs was quantified by calculating the percentage of eccDNA intervals overlapping each feature type. For eccDNAs overlapping multiple features, hierarchical classification prioritized coding regions first, followed by non-coding regions.

### Genome annotation of coding and non-coding elements for human

Genome annotation was performed using HOMER (v4.11) (annotatePeaks.pl) with genome-specific GRCh38.p14.gtf. Annotation was applied separately to CReSIL eccDNA calls, Circle-Map calls, and merged or overlapping sets.

### Repeat annotation for yeast datasets

Repetitive sequences within eccDNA molecules were annotated using RepeatMasker (v4.1.7-p1) with a custom *S. cerevisiae* repeat library (analysis was performed for datasets with accessions: SRR35009460, SRR35009461, SRR35009462, SRR35009463). The custom repeat library was generated by integrating *de novo* repeat discovery and tandem repeat detection. Briefly, a RepeatModeler database was constructed from the *S. cerevisiae* reference genome (GenBank accession R64 with KP263414.1 yeast mitochondrial DNA sequence integrated) using the rmblast alignment engine (v2.14.1). RepeatModeler (v2.0.7) was then employed to identify and classify *de novo* repeat families, including long terminal repeat (LTR) retrotransposons. To capture simple sequence repeats, Tandem Repeat Finder (TRF, v4.09.1; parameters: 2 5 5 80 10 50 2000 -h) was applied to the genome, and repeat sequences were extracted from the TRF output using a custom awk parser, followed by additional filtering with a Python script. The resulting tandem repeat sequences were combined with the RepeatModeler families to produce a unified repeat library. This library was subsequently used as input for RepeatMasker (v4.1.7-p1) to annotate repetitive elements in eccDNA assemblies, employing the parameters -a -s -nolow -html -noisy -nocut. The proportion of eccDNA molecules containing repetitive sequences was quantified, and repeat composition was expressed as the percentage of the total eccDNA sequence length.

### Repeat annotation for human datasets

Repetitive elements in eccDNA assemblies from the human datasets were annotated using RepeatMasker (v4.1.7-p1). RepeatMasker (v4.1.7-p1) was run within the dfam/tetools Docker container using the built-in human repeat library. The software was executed with the parameters -a -s -xsmall -html -noisy -species human, which together enabled alignment-based classification of repeats, increased sensitivity, lowercase masking of repetitive regions, generation of detailed reports in both text and HTML formats, and the application of the curated species-specific human library. The latest Docker image of the dfam/tetools container (bundled with RepeatMasker v4.1.7-p1) was used. For each dataset, the proportion of eccDNA molecules overlapping repetitive elements was calculated, and repeat composition was summarized as the percentage of total eccDNA sequence length attributed to each repeat class (e.g., LINEs, SINEs, LTR retrotransposons, DNA transposons, and simple repeats). Analysis was performed for datasets with accessions: SRR35178673, SRR35178674, SRR35178675, SRR35178676, SRR35178677, SRR35178678.

### HCR IF and HCR-FISH of HEK293T cells

Aliquots (5 ml) of HEK293T cells grown in suspension as described^27^ and stressed with 0.5 mM sodium arsenite (LabChem, Cat# LC228709) for 1 h 20 min were fixed in 3.7% formaldehyde in 1X PBS, pH 7.4 for 10 min at 21 °C. Fixed cells were washed 3 times in 1X PBS and used immediately or stored in 70% (v/v) ethanol at -20 °C. The cells were immobilized on poly-L-Lysine (1 mg/ml; MP Biomedicals, Cat# 102691) covered eight-well chamber cover glass (#1.5 high performance, Cellvis) in 1X PBS for 16-24 h at 4 °C. Consumables for HCR IF and HCR-FISH were ordered from Molecular Instruments, Inc. Lot identifications are provided in the Materials availability section. The design of HCR-FISH probes for AHNAK mRNA and DNA is presented in Figure S3A. The hybridization step with HCR-FISH probes for AHNAK RNA (16-30 nM) or with T30-Q570 (100 nM; Biosearch Technologies, Cat # T30-Quasar 570-1) was as per vendor’s protocol. The hybridization step with HCR-FISH probes for AHNAK, MYC or MAPT DNA targets was performed as follows. First, the DNA-FISH probes (20-30 nM) were applied on slides with immobilized cells in 70% (v/v) formamide (VWR Life Science, Cat# 0606-100ML) in 2 × SSC. The slides were placed on a heat block set at 75 °C and incubated for 3 min; then the slides were quickly transferred to a heat block set at 37 °C and incubated for 15 min. Following hybridization with DNA (40 nM) with or without RNA (30 nM) probes and washes were carried out at 37 °C as per vendor’s protocol. The subsequent immunostaining step was performed using primary antibodies (Table S1) for 1 h 30 min at 21 °C followed by three washes with 1 × PBS with 0.1 % Tween-20, pH 7.4 for 5 min each. Slides were then incubated with corresponding secondary antibodies for 1 h at 21 °C and washed four times with 1X PBS with 0.1 % Tween-20, pH 7.4 for 5 min each. The amplification step was as per vendor’s protocol. Detection of AHNAK mRNA was carried out using amplifier B2-546. Detection of DNA targets was carried out with amplifier B3-647. All washing steps were as per vendor’s instruction for cells on slides. Samples were stained with 0.8 μg/ml Hoechst 33342 (Thermo Scientific, Cat# 62249) in 1X PBS and imaged immediately or covered with ProLong Glass antifade mounting media (Invitrogen, Cat# P36980) and imaged later. Fluorescence images were acquired with a Zeiss LSM880 confocal microscope with a Plan-Apochromat 63×/1.4 Oil DIC M27 objective using Airyscan function with super-resolution mode and 647, 561, 488 and 402 nm laser lines. Approximately 40-50 Z-stacks (0.2 μm depth) were taken from each visual field. Airyscan processing was done by ZEN Black software in 3D mode (automatic setting). Images in figures represent one middle Z-stack; display values were optimized, and signals for proteins and nucleic acids were pseudocolored.

### Design and cloning of CRISPR to target cytoplasmic DNA in yeast

The search for overrepresented elements within DNA from yeast stress granule cores revealed retrotransposons (Ty) to be abundant, with Ty1 dominant. The MEME-1 suite^70^ identified 17-20-mers that were flanked by the ‘NGG’ protospacer adjacent motif. The search (parameters: meme -text -minw 30 -maxw 40 -nmotifs 3) yielded 5’- CCA GGT CAA CCA GGT CTT TAT ATA GAC C<u>AG G</u>AT GAA CTA G-3’ and 5’- CTA GTT CAT CCT GGT CTA TAT AAA GAC C<u>TG G</u>TT GAC CTG G-3’ as most overrepresented motifs for potential targeting Ty11 and Ty12 as 5’- ACC AGG TCT TTA TAT AGA CC-3’ and 5’-TCC TGG TCT ATA TAA AGA CC-3’, respectively. Plasmid pML104^71^ (Addgene #67638; Table S3) carrying Cas9 under the GAP promoter and a nuclear localization signal (NLS) at the C-terminus was used as a backbone. To create a control plasmid, NLS (SV40) was exchanged to a strong nuclear export signal^72^ (NES) by linearization of pML104 with *Mlu*I and *Eag*I, the digest was followed by introduction of the NES-containing insert by Gibson assembly (New England Biolabs, Cat# E5510) using primers NES-For and NES-Rev (Table S4). The constitutive GAP promoter was swapped to the inducible GAL1 promoter by the *EcoR*I, *SgrD*I digest of a plasmid from the previous step (pML104-GAP-Cas9-NES) and subsequent Gibson assembly of the linearized product with the GAL1-encomposing sequence and primers GAL1-For and GAL1-Rev (Table S4). This yielded plasmid pML104-GAL1-Cas9_NES_-Scaffold (Table S3). To eliminate scaffold RNA (tracrRNA) sequence, the plasmid from the previous step was digested with *Bmt*I and *Eag*I, and an insert without scaffold RNA (CON-SNR52; Table S4) was cloned in by Gibson assembly using primers CON-For and CON-Rev, yielding plasmid pML104-GAL1-Cas9_NES_ (Tables S3 and S4). To enhance cytoplasmic cleavage by Cas9, two cassettes for expression of target RNA (crRNA) were cloned into the pML104 backbone. The first targeting RNA (Ty11) complementary to 5’-ACC AGG TCT TTA TAT AGA CC-3’ of Ty1 expressed under SNR52 was inserted using primers Ty11-For and Ty11-Rev (Table S4) with the *Swa*I, *Bcl*I plasmid cleavage and ligation as described^71^. A second cassette targeting another sequence in Ty1 (5’-TCC TGG TCT ATA TAA AGA CC-3’) was designed utilizing the approach as described^73^ where the 3′-end of the gRNA was fused with the self-cleaving HDV ribozyme, which would protect the 5′-end of the RNA from cytoplasmic exonucleases (Ty12^HDV^). To monitor expression of gRNA, the Mango-III(A10U) aptamer^74^ was inserted in lieu of the L4 (UUCG) loop of the HDV ribozyme^75^ generating Ty12-MIII-HDV (Table S4). To insert Ty1-2-MIII-HDV, the pML104 backbone with encoded Ty11 (see above) was digested with *Sph*I and *Eag*I and further assembled with Ty1-2-MIII-HDV using Ty12-For and Ty12-Rev (Table S4). Plasmid where Cas9 was deactivated (dCas9 with D10A and H840A mutations) was generated using QuikChange Lightning site-directed mutagenesis (Agilent Technologies, Cat# 200519) and sense and anti-sense mutagenesis primers a29c and c2518g-a2519c (Table S4). To check expression levels of Cas9 in the cytoplasm, the mCherry-NES fluorescent tag was introduced at the C-terminus of Cas9-NES by linearizing pML104-GAL1-Cas9_NES_-Ty1 (Table S3) with *Mlu*I and *TspM*I and ligating the plasmid with a 1640 bp DNA containing the mCherry tag (pML104-GAL1-Cas9_NES_-mCherry_NES_-Ty1-MIII). The pBY011 plasmids encoding *S. cerevisiae* CHD1 and GCN5 genes under the GAL1/10 promoter was obtained from a DNASU plasmid repository (clone IDs: ScCD00095307 and ScCD00011312, respectively). Alteration of pBY011 and pML104 for introduction of CHD1, GCN5 and additional controls was made by GenScript. Plasmids were sequenced at Plasmidsaurus (www.plasmidsaurus.com) and deposited with Addgene (Table S5).

### Confocal microscopy of yeast transformed with CRISPR targeting cytoplasmic DNA

Yeast transformations were performed using the lithium-acetate protocol^76^ with pretreatment^77^ of cells with DTT to increase transformation efficiency. Transformation mixtures received 1 µg of modified pML104 for expressing CRISPR components. For transient co-expression of CRISPR machinery with CHD1 or GCN5, 0.3 μg of a corresponding plasmid encoding CHD1 or GCN5 was added to the transformation mixtures. Cells carrying plasmid(s) were selected on the SD solid media with kanamycin and 2% (w/v) D-(+)-glucose lacking uracil (SD(-URA)Dex; 30 °C for 3 days). All transformants were re-selected by replating 2-3 medium size colonies on the same SD(-URA)Dex solid media and incubating the plates for two days at 30 °C. Plates (stored at 4 °C) from the previous round were re-streaked once again on the SD(-URA)Dex solid media and incubated for 2-3 days at 30 °C (the third generation). For each transformed variant of YER165W PAB1-GFP, yeast cultures were started at OD_600_ of 6 x 10^-3^ in 10 ml of the SD(-URA)Dex media and grown at 30 °C at 220 rpm until OD_600_ of ∼ 3. The cultures were further diluted in the same media to OD_600_ of 0.1 and continued to grow until OD_600_ of 2-3. The cells were collected by centrifugation at 16000 × *g* for 30 s at 21 °C, dissolved in sterile deionized water, and grown overnight in SD(-URA) containing 1.5% w/v D-(+)-glucose and 0.75% w/v D-(+)-galactose [SD(-URA)Dex-Gal] to express Cas9 (with or without CHD1 or GCN5) moderately. The Cas9 expression was initiated with cultures diluted to 0.15 OD_600_. Experiments with untransformed YER165W Pab1-GFP were carried out in the identical manner except for inclusion of 20 mg/l uracil to the SD media. Overnight cultures were diluted in pre-warmed to 30 °C SD(-URA)Dex-Gal, passed through one doubling time (∼ 4 h) followed by immediate oxidative stress (40 min at 225 rpm) with addition of NaN_3_ directly to the cultures to 0.5% (w/v) final. Cells were fixed with fresh 4.5% (v/v) formaldehyde for 40 min at 21 °C while shaking at 100 rpm, and collected by centrifugation at 4500 × *g* for 10 min. Yeast pellets were washed with ice-cold 100 mM K-phosphate, pH 6.5, incubated on ice for 10 min and re-centrifuged at 4500 × *g*. The washing procedure was repeated twice. Cells were dissolved in 1X PBS, pH 7.4 and analyzed by confocal light microscopy.

### Fluorescence microscopy of fixed yeast cells

Fixed yeast cells were applied on concanavalin A (MP Biomedicals, Cat# 150710) or poly-L-Lysine (MP Biomedicals, Cat# 102691) (for spheroplasts, see below) coated eight-well chamber cover glass (#1.5 high performance, Cellvis, Cat# C8-1.5H-N), then incubated for at least 2-3 h to immobilize cells (or spheroplasts) on slide surfaces. Excess 1X PBS was aspirated, immobilized cells were washed 2 times with 1X PBS, and stained with 1 μg/ml DAPI (Thermo Scientific, Cat# 62248) to visualize DNA and mitochondria. Fluorescence images were acquired with a Zeiss LSM880 confocal microscope (see above) using the super-resolution mode and the line-sequential option. Scans were taken in channel 1 of 488 nm laser, and channel 2 of 405 nm laser. Expression levels of a Cas9 version tagged with mCherry were checked in channel of 561 nm laser. Airyscan Z-stacks were programmed at a step size of 0.14-0.2 μm. Airyscan Z-stacks were processed by Zen Black software in 3D mode (strength parameter set to 5).

### Fluorescence microscopy of yeast spheroplasts

To check expression levels of gRNA, strain JD1370 without any fluorescent fusions was transformed with plasmid pML104-GAL1-Cas9_NES_-Ty1-MIII encoding gRNA-1 and gRNA-2 with the RNA aptamer Mango-III(A10U) (Table S4). Second generation transformants [SD(-URA)Dex] were grown for 24 h to OD_600_ of 5 at 30 °C while shaking at 220 rpm, then diluted and grown in the same non-induction media SD(-URA)Dex to OD_600_ of 0.7-0.8. Cells were collected by centrifugation at 16000 × *g* for 30 s and dissolved either in induction media SD(-URA)Gal with 2% w/v D-(+)-galactose (Figure S4D) or non-induction media SD(-URA)Dex. After overnight growth, cells were diluted in appropriate media, cultured (10 ml) to OD_600_ of 0.45-0.65 and stressed as above with 0.5% (w/v) NaN_3_ for 30 min at 30 °C while shaking at 225 rpm (Figure S4C, left). Untransformed strain JD1370 was used as a control (Figure S4C, right). The control parallels were fermented and carried out in the manner identical to experiments except for inclusion of 20 mg/l uracil to the SD media. Cells were fixed and washed as above. The cell pellets were resuspended in 1 ml of ice-cold 1M Sorbitol in 50 mM potassium phosphate, pH 7.5. To digest cell walls, aliquots of 0.25 ml were supplemented with 2-mecraptoethanol (0.3-0.4 μl of 99% stock) and zymolyase (30 units; ZymoResearch, Cat# E1004), and incubated for 1 h at 30 °C while shaking at 300 rpm. Spheroplasts were collected by centrifugation at 500 × *g* for 6 min at 21 °C, then dissolved in buffer A [30 mM potassium phosphate, pH 7.5 (at 20 °C), 140 mM KCl, 1.5 mM MgCl_2_, 1M Sorbitol] and centrifuged at 500 × *g* for 10 min at 21 °C. The washing procedure was repeated. Finally, spheroplasts were gently dissolved in 0.2 ml of buffer A and stored on ice (up to 2 days). Prior to imaging, spheroplasts were immobilized on poly-L-Lysine coated slides, then biotinylated TO1 fluorophore^74^ was added to 200 nM and fluorescent micrographs were immediately acquired using LSM 880 with Airyscan (laser: 488 nm; see above).

### Post-stress recovery of CRISPR transformant assays

YER165W PAB1-GFP strain was transformed with CRISPR variants as specified in Figure 5A. The initiator cultures were grown in SD(-URA)Dex at 30 °C at 220 rpm until OD_600_ of 0.5-1.0. The cultures were diluted in the same media to OD_600_ of ∼ 0.1 and continued to grow until OD_600_ of 1.0. The cells were then diluted in expression media (SD(-URA)Dex-Gal) to OD_600_ of ∼ 0.04, grown overnight and stressed as above (40 min at 225 rpm) at OD_600_ of 0.4-0.5. After pelleting at 16000 × *g* for 1 min at 21 °C, the cells were washed in ultra-pure water, collected again and dissolved in water to bring OD_600_ to 1.0. Recovery was assayed on solid expression media (SD(-URA)Dex-Gal) by spotting 3 μl of ten-fold dilutions (Figure 5A) and incubating plates at 30 °C for 4 days. The recovery efficiency was quantified (Figure 5B) using protocol as described^78^ and one-way ANOVA test followed by Tukey’s test (*df* between groups 4; *df* within group 20). The same stressed cells were also cultured in liquid expression media using OD_600_ of 0.01 to initiate the growth. The growth was monitored over 2 days at 30 °C at 220 rpm (Figure 5C).

## DATA AND MATERIALS AVAILABILITY

Raw sequencing data have been deposited in the Sequence Read Archive (Table S2). Sequencing data for yeast and human DNA are available under BioProjects PRJNA1306188 and PRJNA1305524, respectively. Plasmids generated in this study are available from the authors under a material transfer agreement with Addgene. Deposited plasmid IDs are listed in Table S5. Requests should be submitted by email to A.R.F.-D. Per manufacturer’s policies, FISH probes can be requested from Molecular Instruments, Inc. referring to lots RTG842 and RTG843 for AHNAK RNA and DNA targets, respectively; lots RTG845 and RTK096 for MYC and MAPT DNA targets, respectively.

