## Supplemental Data (figures and tables) for "Cytoplasmic circular dsDNA is a key constituent of stress granules"

### Supplementary Figures

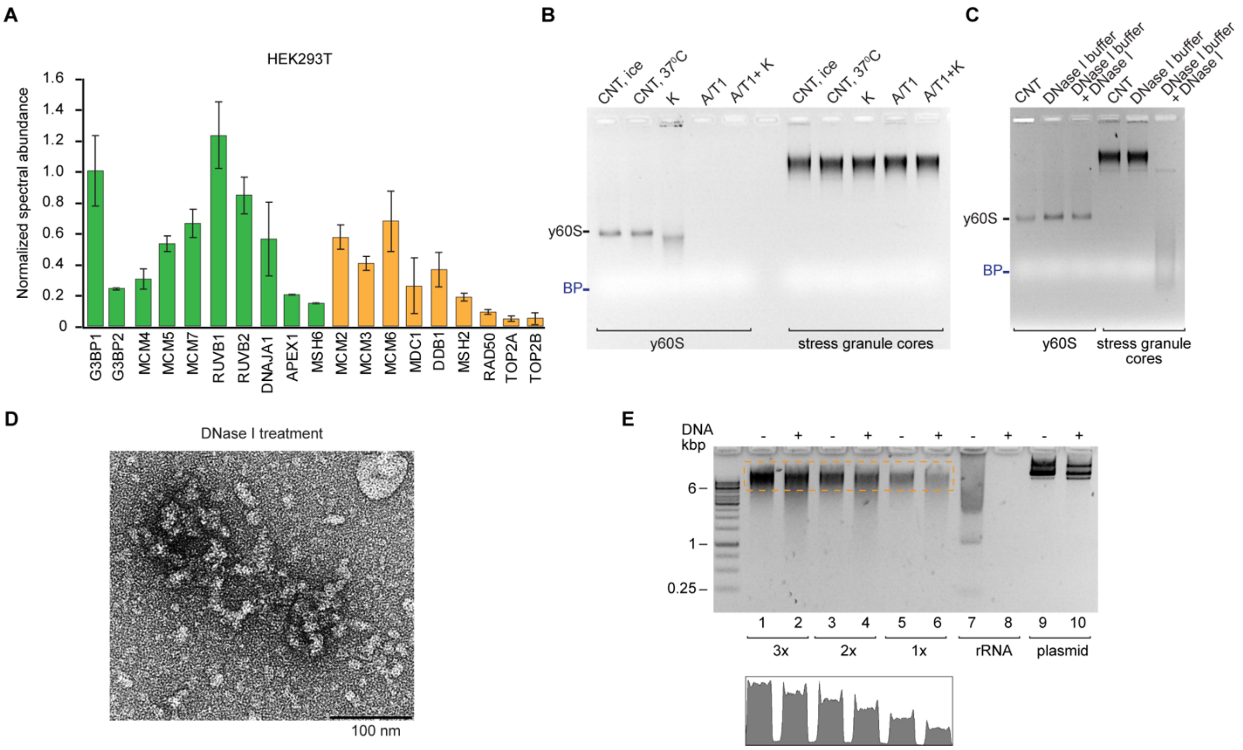


**Figure S1. Proteinaceous and DNA constituents of stress garnule cores,** related to Figure 1.

(A) Abundance of representative DNA-binding proteins in the proteomes of stress granule cores isolated from early-log HEK293T cells[1]. Green color denotes DNA-binders also found in immunoprecipitated stress granules from late-log U-2 OS cells[2]; orange color marks DNA-binders additionally identified in the cores of HEK293T cells. Relative abundance of proteins between datasets was normalized using G3BP1 (data are mean ± s.d. from $n = 3$ biological replicates).

(B) Native agarose gel electrophoresis (ethidium bromide stain) of the yeast large ribosomal subunit (y60S) and yeast stress granule cores treated with proteinase K, ribonucleases A and T1 (A/T1), and combination of these enzymes. Incubation was performed at 37 °C for 24 hours.

(C) Native agarose gel electrophoresis (ethidium bromide stain) of y60S and yeast stress granule cores treated with DNase I. Incubation was performed at 37 °C for 1 hour.

(D) Negative-stain electron micrograph of yeast stress granule cores treated with DNase I for 30 minutes at 30 °C. Magnification (× 13,000).

(E) Alkaline hydrolysis of total nucleic acids extracted from yeast stress granule cores (15, 10 and 5 ng), rRNA (15 ng) and 11 kbp plasmid (15 ng), analyzed by non-denaturing agarose gel electrophoresis (SYBR Gold stain). Samples marked (+) were incubated with 50 mM KOH for 15 min at 95 °C. Bottom graph, integrated signal for the dashed box. In (B) and (C), BP stands for bromophenol blue.

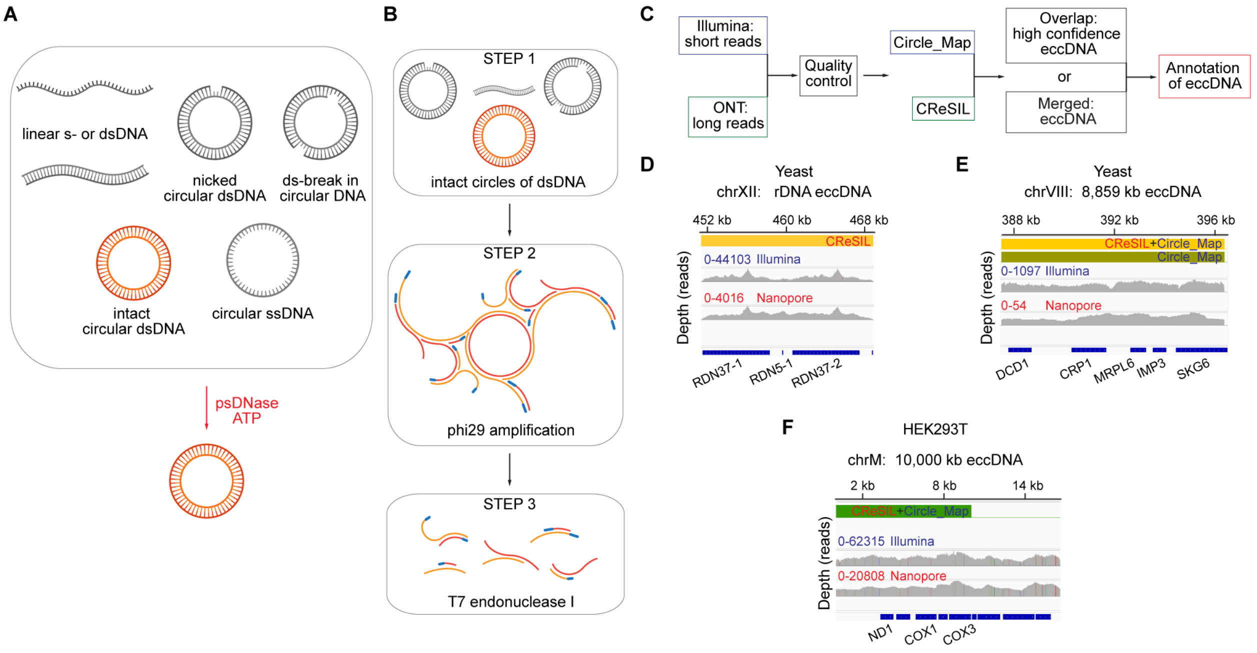


**Figure S2. Workflow for analysis of DNA from stress granule cores by eccDNA assemblers,** related to Figure 2.

(A) Substrate specificity of psDNase. Species of DNA hydrolyzed by psDNase in the presence of ATP are in black.

(B) Main steps of modified Circle-Seq treatment applied to enrich (step 1), amplify (step 2) and debranch (step 3) circular DNA from stress granule cores for further sequencing.

(C) Bioinformatic workflow of processing DNA reads for assembly into eccDNA circles.

(D-F) Examples of genomic regions covered by short (Illumina) and long (Nanopore; ONT) reads, whose sequences are assembled in eccDNAs for yeast (D) and (E) and human (F) stress granule cores. Panels (A) and (B) created with BioRender.com.

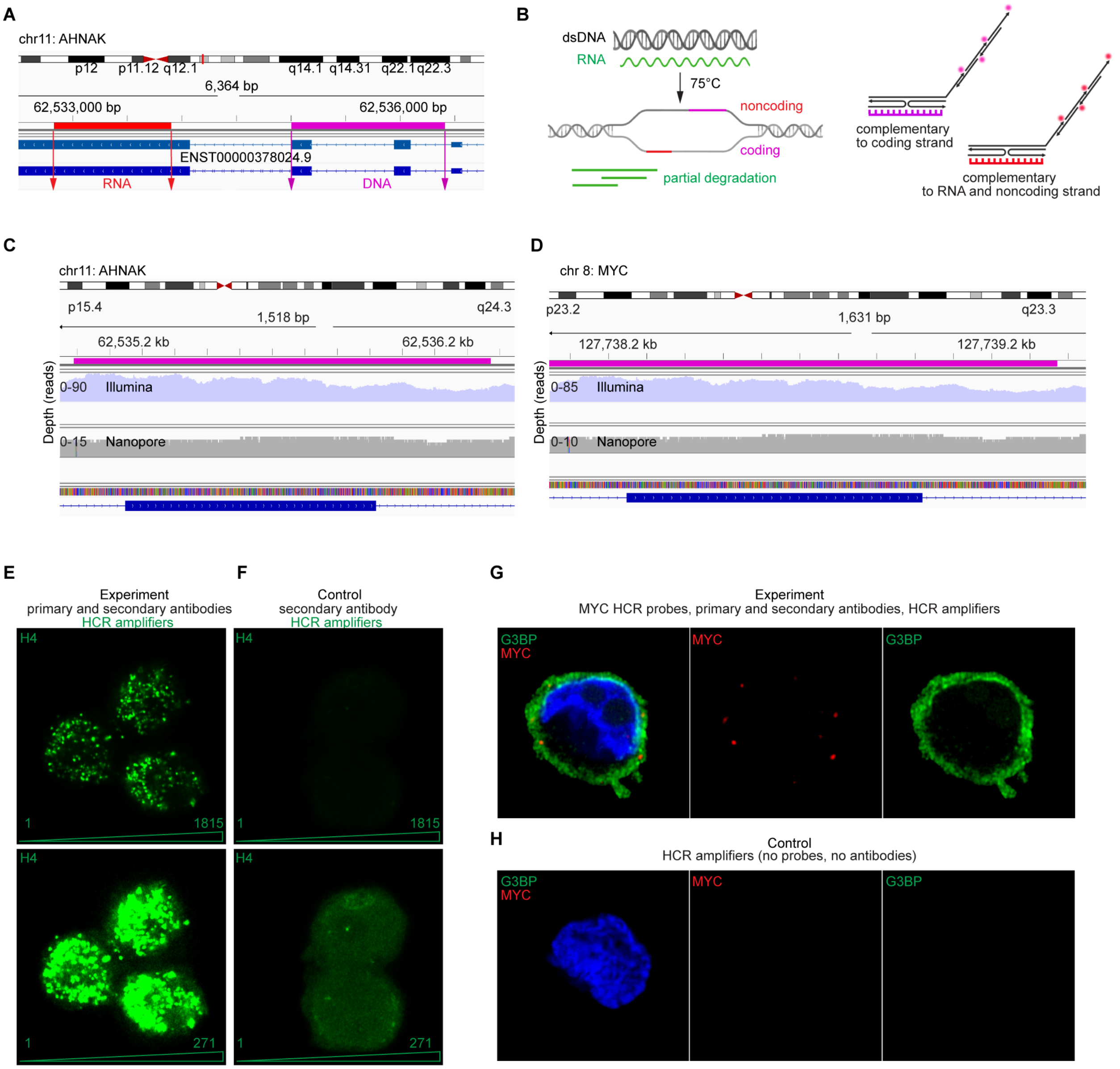


**Figure S3. Detection of stress granule core constituents in HEK293T cells by HCR fluorescence,** related to Figure 3.

(A) Exemplary design of FISH probes for targeting DNA (magenta) and RNA (red) counterparts of a gene of interest.

(B) Schematic of heat-denaturation step of HCR-FISH protocol aimed to detect DNA coding region (magenta). Concurrent detection of transcribed RNA with corresponding region of non-coding DNA strand is indicated (red).

(C-D) Representative DNA Illumina and Nanopore reads covering regions within AHNAK (C) and MYC (D) genes to which HCR-FISH probes (magenta) were designed.

(E-F) Abundance of histone H4 in the cytoplasm of HEK293T cells. Two gains (top and bottom) demonstrate sensitivity and specificity of HCR immunolabeling.

(G-H) HCR amplifiers are highly specific to primary targets (FISH probes or antibodies).
In (E-H) all images represent one middle Airyscan confocal Z-stack of 0.2 μm. Nuclei were stained by Hoechst 33342 (blue) in (G) and (H).

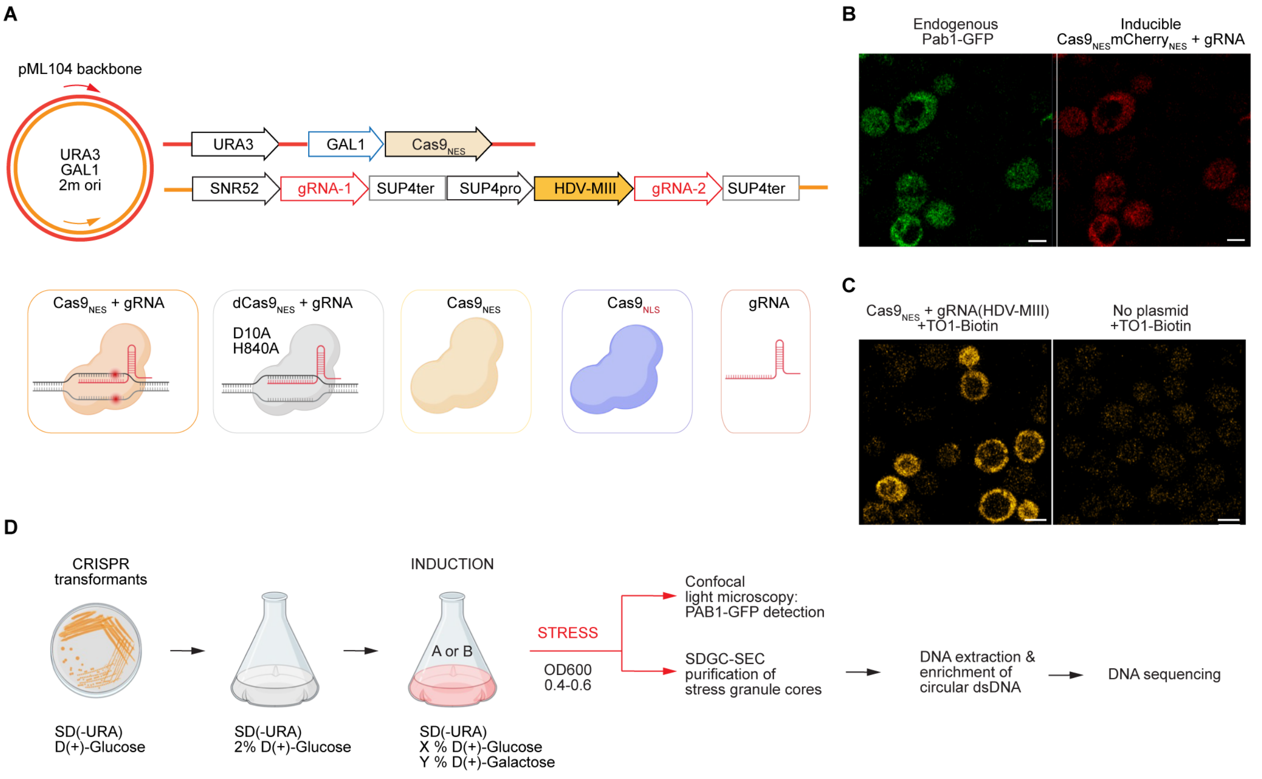


**Figure S4. Design of CRISPR experiments for targeting cytoplasmic eccDNA in yeast stress granule cores,** related to Figure 4.

(A) Major regulatory and coding motifs of CRISPR elements in modified vector pML104. Botton panel displays experimental variants of yeast transformants with CRISPR-pML104. NES, nuclear export signal; NLS, nuclear localization signal.

(B) Transiently expressed Cas9$_{NES}$mCherry$_{NES}$ localizes to cytosol together with PAB1-GFP (condition A, see (D), no stress).

(C) Detection of gRNA in yeast cytoplasm using fluorophore TO1-Biotin and fluorogenic RNA aptamer Mango-III(A10U) (see Methods) encoded in HDV ribozyme at 5'-end of gRNA-2 (see (A)). The experiments were conducted with strain JD1370 free from endogenous or transient fluorescent fusions.

(D) Growth steps of CRISPR-pML104 transformants with indication of major alterations in media, viz. changing D(+)-glucose to D(+)-galactose ratio to induce expression of Cas9 and addition of sodium azide to stress cells. Condition A: 0.2% w/v D-(+)-glucose and 2% w/v D-(+)-galactose for 4 h. Condition B: 1.5% w/v D-(+)-glucose and 0.75% w/v D-(+)-galactose, overnight.

In (B) and (C), Airyscan images of middle Z-stack plane (0.14 μm). Scale bar 2 μm. Panels (A) and (D) created with BioRender.com.

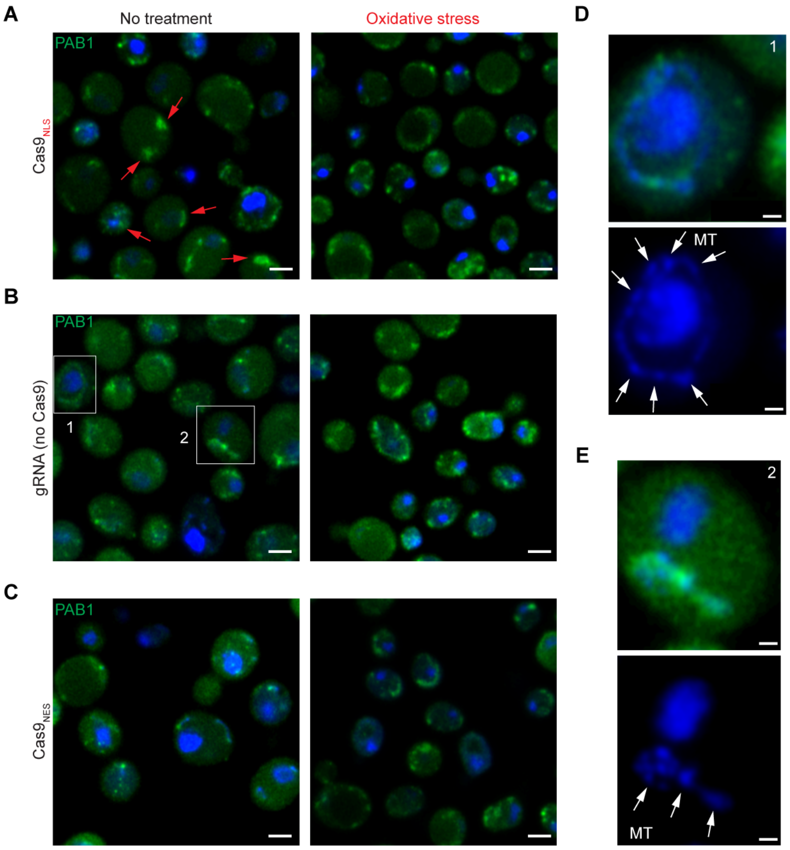


**Figure S5. Phenotypic changes of *S. cerevisiae* in response to transient expression of individual CRISPR elements,** related to Figures 4 and 5.

(A-C) Confocal microscopy of *S. cerevisiae* with endogenous PAB1-GFP (green) and transiently expressed Cas9 (GAL promoter) or gRNAs (see Figure S4A). Treatment conditions and nuclear (NLS) or cytoplasmic (NES) tags are indicated. In (A), arrows (red) indicate accumulation of PAB1-GFP signal observed in the absence of stress.

(D-E) Colocalization of PAB1-GFP with mitochondria in the absence of stress (framed in (B)). Upper images (488 and 405 nm channels) are split into individual channels to mark mitochondria (lower panels; arrows).

All images represent one middle Airyscan confocal Z-stack of 0.14 µm with nuclei and mitochondria stained by DAPI (blue). Scale bars 2 µm in (A-C) and 0.5 µm in (D) and (E).

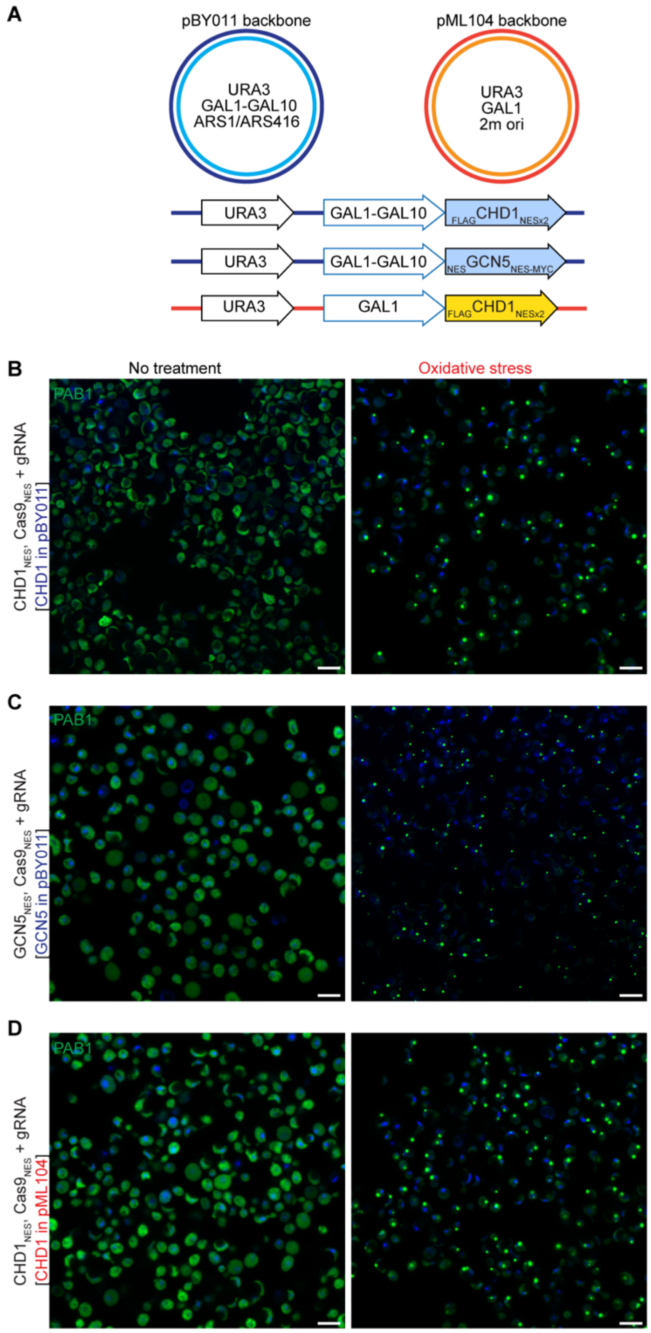


**Figure S6. Chromatin binders CHD1 and GCN5 compete out CRISPR and restore stress granule formation in cytoplasm,** related to Figures 4 and 5.

(A) Schematics of pBY011 and pML104 main elements used to express chromatin remodeler CHD1 or histone acetyltransferase GCN5.

(B) Transient co-expression of cytoplasmic CHD1 encoded into pBY011 backbone with cytoplasmic CRISPR preserves canonical stress response.

(C) Transient co-expression of cytoplasmic GCN5 encoded into pBY011 backbone with cytoplasmic CRISPR preserves canonical stress response.

(D) Transient co-expression of cytoplasmic CHD1encoded into pML104 backbone with cytoplasmic CRISPR preserves canonical stress response.

Right panels in (B) to (D), cells treated with 0.5% w/v sodium azide (Methods). All images represent one middle Airyscan confocal Z-stack of 0.14-0.17 μm with nuclei and mitochondria stained by DAPI (blue). Scale bar 5 μm.

### Supplementary Tables

**Table S1. Antibodies used in IF and HCR IF light microscopy imaging of HEK293T cells.**

| Primary Antibody [dilution] | Secondary Antibody [dilution] | Amplifier |
|---|---|---|
| Anti-VCP, mouse, monoclonal [1:100] (Santa Cruz Biotechnology, Cat# sc-57492) | Anti-mouse, donkey [1:500], B5 initiator Molecular Instruments, Inc. | B5-647 |
| Anti-histone H4, rabbit, monoclonal [1:70 – 1:100] (Abcam, Cat# ab177840) | Anti-rabbit, donkey [1:500], B1 initiator Molecular Instruments, Inc. | B1-488 |
| Anti-histone H3.1, rabbit, monoclonal [1:30 to 1:90] (Novus Biologicals, Cat# NBP3-26228) | Anti-rabbit, donkey [1:500], B1 initiator Molecular Instruments, Inc. | B1-488 |
| Anti-G3BP, mouse, monoclonal [1:200] (Abcam, Cat# ab56574) | Anti-mouse, donkey [1:500], B5 initiator Molecular Instruments, Inc. | B5-647 |
| Anti Lamin A + Lamin C, rabbit, monoclonal [1:400] (Abcam, Cat# ab108595) | Anti-rabbit, donkey [1:800], B1 initiator Molecular Instruments, Inc. | B1-488 |
| Anti-Caprin1, rabbit, polyclonal [1:400] (Abcam, Cat# ab244360) | Anti-rabbit, donkey [1:750], B4 initiator Molecular Instruments, Inc. | B4-546 |
| Anti-TOMM20, rabbit, monoclonal [1:250] (Abcam, Cat# ab186735) | Anti-rabbit, donkey [1:500], B1 initiator Molecular Instruments, Inc. | B1-488 |
| Anti-DNA, mouse, monoclonal [1:200] (Thermo Fisher Scientific, Cat # 690014S) | Anti-mouse, goat [1:800], Alexa Fluor 568 (Invitrogen, Cat# A11004) | N/A |

**Table S2. Sequence Read Archive deposition summary for raw DNA sequences.**

| Bioproject | Accession | Species | Sequencing platform | Sample name |
|---|---|---|---|---|
| PRJNA1306188 | SRR34996475 | *S. cerevisiae* | Illumina | Cas9nes_gRNA_01 |
| PRJNA1306188 | SRR34996474 | *S. cerevisiae* | Illumina | Cas9nes_gRNA_02 |
| PRJNA1306188 | SRR34996473 | *S. cerevisiae* | Illumina | dCas9nes_gRNA_01 |
| PRJNA1306188 | SRR34996472 | *S. cerevisiae* | Illumina | dCas9nes_gRNA_02 |
| PRJNA1306188 | SRR34996471 | *S. cerevisiae* | Illumina | No_tranform_01 |
| PRJNA1306188 | SRR34996470 | *S. cerevisiae* | Illumina | No_tranform_02 |
| PRJNA1306188 | SRR34996469 | *S. cerevisiae* | Illumina | Cas9nes_01 |
| PRJNA1306188 | SRR34996468 | *S. cerevisiae* | Illumina | Cas9nes_02 |
| PRJNA1306188 | SRR35009179 | *S. cerevisiae* | Illumina | Cas9nes_gRNA_CHD1nes_01 |
| PRJNA1306188 | SRR35009178 | *S. cerevisiae* | Illumina | Cas9nes_gRNA_CHD1nes_02 |
| PRJNA1306188 | SRR35009463 | *S. cerevisiae* | Illumina | ySCGs_Illumina_01 |
| PRJNA1306188 | SRR35009462 | *S. cerevisiae* | Illumina | ySCGs_Illumina_02 |
| PRJNA1306188 | SRR35009461 | *S. cerevisiae* | ONT | ySGCs_ONT_01 |
| PRJNA1306188 | SRR35009460 | *S. cerevisiae* | ONT | ySGCs_ONT_02 |
| PRJNA1305524 | SRR35178678 | *H. sapiens* | ONT | 7E_Dna_nanopore |
| PRJNA1305524 | SRR35178677 | *H. sapiens* | ONT | 8E_Dna_nanopore |
| PRJNA1305524 | SRR35178676 | *H. sapiens* | ONT | 9E_Dna_nanopore |
| PRJNA1305524 | SRR35178675 | *H. sapiens* | Illumina | 7E_Dna_Illumina |
| PRJNA1305524 | SRR35178674 | *H. sapiens* | Illumina | 8E_Dna_Illumina |
| PRJNA1305524 | SRR35178673 | *H. sapiens* | Illumina | 9E_Dna_Illumina |
| PRJNA1305524 | SRR34981497 | *H. sapiens* | ONT | hSGCs 1_early_ONT |
| PRJNA1305524 | SRR34981496 | *H. sapiens* | ONT | hSGCs 2_late_ONT |
| PRJNA1305524 | SRR34981495 | *H. sapiens* | ONT | hSGCs 3_early_ONT |
| PRJNA1305524 | SRR34981494 | *H. sapiens* | ONT | hSGCs 4_late_ONT |
| PRJNA1305524 | SRR34981493 | *H. sapiens* | ONT | hSGCs 5_early_ONT |
| PRJNA1305524 | SRR34981492 | *H. sapiens* | ONT | hSGCs 6_late_ONT |
| PRJNA1305524 | SRR35328255 | *H. sapiens* | Illumina | hSGCs 1_early_Illumina |
| PRJNA1305524 | SRR35328254 | *H. sapiens* | Illumina | hSGCs 2_late_Illumina |
| PRJNA1305524 | SRR35328253 | *H. sapiens* | Illumina | hSGCs 3_early_Illumina |
| PRJNA1305524 | SRR35328252 | *H. sapiens* | Illumina | hSGCs 4_late_Illumina |
| PRJNA1305524 | SRR35328251 | *H. sapiens* | Illumina | hSGCs 5_early_Illumina |
| PRJNA1305524 | SRR35328250 | *H. sapiens* | Illumina | hSGCs 6_late_Illumina |

**Table S3. Cloning materials for CRISPR targeting cytoplasmic eccDNA with Ty1 in yeast.**

| Plasmid Name | Description |
|---|---|
| pML104 | Original (Addgene #67638)<br>GAP-$Cas9_{NLS}$<br>Scaffold (tracrRNA) under SNR52<br>No crRNA |
| pML104-GAL1-$Cas9_{NES}$-Scaffold | GAL1-$Cas9_{NES}$<br>Scaffold RNA (tracrRNA) / SNR52<br>No crRNA |
| pML104-GAL1-$Cas9_{NES}$ | GAL1-Cas9-NES<br>No scaffold RNA<br>No crRNA |
| pML104-GAL1-$Cas9_{NES}$-Ty1 | GAL1-Cas9-NES-Ty1<br>crRNA-1: SNR52-Ty11<br>crRNA-2: SUP4-$Ty12^{HDV}$ (minus MIII insert) |
| pML104-GAL1-$Cas9_{NES}$-Ty1-MIII | GAL1-Cas9-NES-Ty1<br>crRNA-1: SNR52-Ty11<br>crRNA-2: SUP4-$Ty12^{HDV\text{-}MIII}$ (plus MIII insert) |
| pML104-GAL1-$dCas9_{NES}$-Ty1 | GAL1-dCas9-NES-Ty1<br>crRNA-1: SNR52-Ty11<br>crRNA-2: SUP4-$Ty12^{HDV}$ |
| pML104-GAL1-$Cas9_{NES}$-$mCherry_{NES}$-Ty1-MIII | GAL1-Cas9-NES-mCherry-NES-Ty1<br>crRNA-1: SNR52-Ty11<br>crRNA-2: SUP4-$Ty12^{HDV\text{-}MIII}$ (plus MIII insert) |

**Table S4. Primers and fragments designed for modification of the pML104 backbone.**

| Name | Sequence |
|---|---|
| NES-For | 5’- cagatttccgagttttctaaacgcgtcattctcgctgat |
| NES-Rev | 5’- tgccaatctaaacgataccacggccgctctagagaaatgg |
| GAL1-For | 5’- gtatcgataagcttgatatcgaattcagtctactagtcagttcgagtttacg |
| GAL1-Rev | 5’- tttctcgtgataggccacctcgtcgacgatattaccgaagatgg |
| CON-SNR52 | 5’- ggccgtggtatcgtttagattggcaattacagtgtcttagctcacatgcttataactaattacatgactcgaagacataaaaaacaaaaaaagatcatttatctttcactgcggagaagtttcgaacgccgaaacatgcgcaccaactttcacttctacagcgtttgaccaaaatcttttgaacagaacattgtagggtgtgaaaaaatgcgcacctttaccg |
| CON-For | 5’- cgagcaaatgcctgcaaatcgctccccatttctctagagcggccgtggtatcgtttagattgg |
| CON-Rev | 5’- gaagtacaactctagattttgtagtgccctcttgggctagcggtaaaggtgcgcatttttcac |
| MIII(U10A) | 5’-GTACGAAGGAAGGTTTGGTATGGGGTAGTTGTCGTAC-3’ |
| Ty1-2-MIII-HDV | 5’- agacataaaaaacaaaaaaagcaccgactcggtgccactttttcaagttgataacggactagccttattttaacttgctatttctagctctaaaacGGTCTTTATATAGACCAGGAaaa**gtcccattcgccacc**GTACGACAACTACCCCATACCAAACCTTCCTTCGTAC**ggtgttgcccagccggcgccagcgaggaggctgggaccatgccgg**ccatctctcccgggggcgagtcgaacgcccgatctcaagatttcgtagtgataaattacagtcttgcgccttaaaccaacttggctaccgagagtatttaattgttgaagaaagagtatactacataacacatata -3’ |
| Ty12-For | 5’- gcatgaggtcgctcttattgaccacacctctaccggcatgagacataaaaaacaaaaaaagcaccgact |
| Ty11-For | 5’- gatcg**accaggtctttatatagacc**-3’ |
| Ty11-Rev | 5’- **ggtctatataaagacctggt**c-3’ |
| Ty12-Rev | 5’- tgagctaagacactgtaattgccaatctaaacgataccactatatgtgttatgtagtatactctttcttcaacaattaaatactctcg |
| a29c-s | 5'-gtattctatcggactggccatcgggactaatagcg-3' |
| a29c-as | 5'-cgctattagtcccgatggccagtccgatagaatac-3' |
| c2518g-a2519c-s | 5'-ctctgagggacgatggcgtccacgtcgtagtc-3' |
| c2518g-a2519c-as | 5'-gactacgacgtggacgccatcgtccctcagag-3' |

**Table S5. Addgene deposit IDs for plasmids used in the study.**

| Name | Sequence ID [deposit] |
|---|---|
| pML104_Gal1_Cas9NES_Ty1MIIIA10U | 215655 [83841] |
| pML104_Gal1Cas9NES_Ty1 | 215656 [83841] |
| pML104_Gal1dCas9NES_Ty1 | 215658 [83841] |
| pML104-Gal1Cas9NES | 215659 [83841] |
| pBY011_Flag_CHD1_NES2 | 215660 [83841] |
| pML104_Ty1_11_12 | 246009 [86470] |
| pML104_Gal1_Cas9NLS | 246012 [86470] |
| pML104_Gal1_Flag_CHD1NES2 | 246015 [86470] |
| pBY011_NES_GCN5_NES_MYC | 246016 [86470] |
